# Leaf microbiome assembly is linked to plant phylogeny

**DOI:** 10.64898/2025.12.17.695039

**Authors:** Pankaj K. Singh, Catarina S. C. Martins, Ramesha H. Jayaramaiah, Eleonora Egidi, Catriona Macdonald, Juntao Wang, Chao Xiong, Bruna Batista, Galaxy Qui, Peter B. Reich, Manuel Delgado-Baquerizo, Brajesh K Singh

**Author notes:** Correspondence: Brajesh K Singh and Pankaj K Singh and.

## Abstract

**Background and Aims:** The plant microbiome is considered as an extended part of the plant genome, and it provides key functions in regulating plant fitness, and stress tolerance. Plants and associated microbiomes have co-evolved over millennia, yet evidence for a strong influence of plant phylogeny in influencing their microbiomes is largely lacking. Our main aims was to identify key drivers of plant microbiome assembly.

**Methods:** Here, we conducted a full factorial experiment that included three levels of soil microbial diversity, five plant species from three functional groups (C3, C4, and C3 nitrogen-fixing), and two moisture availability levels.

**Results:** Our results showed that host identity and plant functional group exerted the strongest effect on leaf microbial assembly, while root and soil microbiomes showed less sensitivity to host selection. The initial soil microbial diversity and community structure significantly impacted soil and root microbial composition, but not leaf microbiomes. Importantly, we observed significant positive linkage between host phylogeny distance and Bray-Curtis dissimilarity index in leaf microbiomes. This finding was further validated through analysis of microbiome data from seven plant species grown across different field and environmental conditions. Interestingly, there was no significant impact of short-term water stress on plant microbial communities.

**Conclusions:** By providing empirical evidence for important role of host selection in shaping plant microbiomes, this study advances our fundamental knowledge of plant-microbe interactions and their co-evolutionary relationships, and enhances our ability to develop future tools harnessing plant microbiome to improve plant health and productivity.

## Introduction

The plant microbiomes, including that in the rhizosphere, roots and leaves, are known to play critical roles in regulating plant fitness and productivity, providing the host with nutrients and resilience against biotic and abiotic stresses in exchange for energy and habitat provision (Trivedi et al. 2020; Li et al. 2025). Some studies reported the importance of host related factors in plant microbiome assembly for Arabidopsis, tomato, rice and maize (Bodenhausen et al. 2013; Bulgarelli et al. 2013; Bai et al. 2015; Edwards et al. 2015). Given that most of the plant microbiota are recruited from the soil (Xiong, et al. 2021a; Xiong, et al. 2021b; Singh et al. 2023b), it is likely that the plant microbiome assembly is determined by the initial soil microbial diversity and its composition (Compant et al. 2019; Trivedi et al. 2020). However, empirical evidence for relative contributions of soil microbiome and host selection is lacking. Further, microbial communities occupying different plant niches (e.g. root, leaf) vary and likely to respond differently to external stimuli. This is because different plant tissues have different physiological and chemical response to external stimuli (Laforest-Lapointe et al. 2016; Morella et al. 2020; Trivedi et al. 2020; Singh et al. 2023a) that, in turn, is likely to influence microbiome assembly. However, such ecological factors are often studied independently, and the relative contributions of host types and niches, soil microbial diversity, and environmental stresses on plant microbiome assembly remain poorly understood (Fitzpatrick et al. 2018; Compant et al. 2019; Xiong, Zhu, et al. 2021; Xiong, Singh, et al. 2021). These knowledge gaps constrain advancement in the fundamental knowledge on plant-microbial co-evolutions and limit our ability to harness host microbiome to improve plant health and productivity.

In addition to the soil microbiome, the role of plant evolutionary processes in determining host microbiome assembly remain largely unknown. For example, current evidence indicates that microbes recognise chemical signals produced by the plant, while the plant immune response ensures only recognised microbes can colonise, corroborating the notion that the microbiome assembly is supported by mutual recognition processes that are co-evolutionary in nature (Fitzpatrick et al. 2020; Trivedi et al. 2020; Singh et al. 2025). Further support is provided by symbiotic relationships, such as those between rhizobia and legumes, as well as mycorrhizal fungi and higher plants (Genre et al. 2020). Similarly, the existence of plant core microbiomes has been proposed as an example of this co-evolutionary process (Toju et al. 2018; Hamonts et al. 2018). Under the co-evolutionary theory, a correlation between plant phylogeny and microbiome composition is expected. This is because plant species that are closely related are more likely to harbour similar microbiomes due to shared evolutionary history and similarities in their internal chemical and physical environments (Heckman et al. 2001). Yet, detailed evidence on whether and to what extent plant phylogenetic relatedness and functional traits reflect the plant microbiome composition remains unclear. Addressing these knowledge gaps are critical to improve mechanistic understanding of plant microbiome assembly and to potentially harness them to increase primary productivity.

In this study, our aim was to determine role and relative contributions of soil microbial diversity, host phylogenetic relatedness, host niches and environmental disturbance (drought-a short-term water stress) in shaping the microbiome of different soil and plant niches across multiple plant species, and functional groups. A soil microbial diversity gradient was employed to test the impact of soil microbial diversity and composition on the root and leaf microbiomes of five plant species belonging to three groups (C3, C4, and nitrogen-fixing plants) from different taxonomic and functional groups. Specifically, our objectives were to (i) assess whether initial soil microbial diversity and composition, or plant species identity is the dominant driver of plant microbiome assembly, and how these factors interact with environmental stressor (drought) to impact plant microbiome assembly, and (ii) evaluate whether host phylogeny was linked to the plant microbiome assembly. We hypothesized that the phylogenetic distance between hosts will explain the variation in their microbiomes. Further, we hypothesized that leaf microbiome will be less affected compared to root microbiomes by disturbance or soil microbial diversity gradient because of progressive host filtering and unique internal physico-chemical properties.

## Material and Methods

We employed combined approach of glasshouse experiments and a field survey to achieve our aims. **Glass house experiment**

### 2.1.1 Initial microbial diversity manipulation, plant establishment and water stress treatment

The experimental design consisted of three levels of soil microbial diversity, five plant species and a disturbance treatment in the form of water stress (drought) followed by a recovery period. The experiment was set up in 2018 using topsoil (0-15 cm) collected from the Hawkesbury Forest Experiment site, Richmond, New South Wales, Australia. The soil (loamy texture, total carbon = 1.65%, total nitrogen = 0.15%, extractable nitrate = 0.6 mg kg^-1^, extractable ammonia = 2.3 mg kg^-1^, extractable phosphorus = 20 mg kg^-1^, pH=6.43, carbon/nitrogen ratio of = 11:1 and volumetric WHC of 15-20%) was sieved through a 5 mm mesh and homogenized into 108 aliquots of 1.9 kg and stored in plastic bags.

We used the dilution-to-extinction approach to create a gradient of soil microbial diversity. We used this approach to obtain different soil biodiversity and composition, rather than using different soils with natural variation in microbial diversity. This is because soils from different location have different abiotic properties (e.g. pH, nutrients, structure) which may affect both microbiomes and plant physiology and thus plant-microbiome interactions. This makes difficult to disentangle effects of soil microbiomes from soil abiotic properties on plant microbiome assembly and interactions. The dilution-to extinction approach overcomes these shortcomings in plant-microbiome research by allowing to keep the soil abiotic properties relatively constant, while the only characteristic change is soil microbial diversity and composition. To obtain the diversity gradient, bagged soil was sterilized using gamma radiation (50kGy each) at ANSTO Life Sciences facilities, Sydney, New South Wales, Australia. The sterilized soil was used for potting, while a subset of unsterilized soil was used to prepare three microbial inocula using a dilution to extinction approach (Trivedi et al. 2019; Martins et al. 2024). Briefly, unsterilized soil was used to prepare a 10-fold serial dilution to create a microbial diversity gradient. The gradient consisted of three dilutions, U (undiluted/10^1^), 10^-2^, and 10^-6^. The inocula were then added in ratio of 1:9 (inocula : soil) to six replicates of previously sterilized soils and incubated for 18 weeks to allow microbial biomass recovery (Supplementary Figure 1).

Our experiment included monocultures of five plant species from three functional groups (C3, C4, and nitrogen-fixing) for the plant treatment. C3 plants were represented by ryegrass (*Lolium perenne*) and phalaris (*Phalaris aquatica*), C4 plants were represented by Rhodes grass (*Choris gayana*) and digitaria (*Digitaria eriantha*), whereas lucerne (*Medicago sativa*) was selected as a nitrogen fixing plant. Initially, seeds were sterilized using a 10% bleach solution for 5 mins and were washed repeatedly with sterile milliQ water. Seeds were then immersed in a composite solution representing individual soil microbial diversity levels and were kept in a growth chamber (25^º^C/15ºC day/night; 14h/10h day/night cycle) for germination on sterile vermiculite. Post germination, 3 cm long seedlings were then transferred to their designated pots. Each pot had six plant and had six replicates for each of the dilution gradients. (Supplementary Figure 1).

Our experiment also included two levels of water stress (drought vs. control) to investigate impact of drought stress on microbiome assembly. Once the plants were established up to 3 cm in height, the pots were divided into two groups with three replicates each and were watered for first 12 weeks at 60% water holding capacity (WHC). For the next two weeks, one group was kept at 60% WHC and was considered as a control, while the other group was kept at 30% of WHC and considered as a drought treatment (Suppl. Fig 1). After two weeks, the drought-treated pots were returned to 60% WHC for three weeks to allow post-drought recovery. After recovery, three soil cores of 15 cm depth were collected from each pot and then combined to obtain a composite sample for each pot. The soil was sieved (2 mm) and stored at -20^º^C. Three leaves per plant were selected for leaf sampling and along with roots were harvested and stored at -20^º^C.

### Surface sterilization, DNA isolation, and amplicon sequencing

After harvesting, root and leaf samples were surface sterilized with 70% ethanol for 2 minutes, followed by 2% sodium hypochlorite for 3 mins, and again 70 % ethanol for 1 min to remove epiphytes. The samples were then washed at least twice with sterile distilled water to remove any remnants of sterilizing agents. After surface sterilization, the leaf and root samples were cut into small pieces ∼2 mm × 2 mm (Qiu et al. 2020). We used 0.25 g of leaf, root, and soil samples for DNA extraction, bead beating was performed and following that DNA was extraction was done using the PowerSoil® DNA Isolation kit (Qiagen Inc., Valencia, CA, USA), following the manufacturer’s instructions. The quality of DNA was determined using the NanoDrop spectrophotometer (NanoDrop Technologies, Wilmington, DE, USA). Microbial community profiling was performed using amplicon-based sequencing methods using an Illumina sequencing platform. The V5-V6 region of bacterial 16S rRNA gene was amplified using 799F and 1115R set of primers (Kembel et al. 2014), and fungal Internal Transcribed Spacer 2 (ITS2) region was amplified using fITS7 and ITS4 primers (Ihrmark et al. 2012)). Sequencing was performed at the Next Generation Sequencing Facility of Western Sydney University (Richmond, New South Wales, Australia).

### Bioinformatics analysis

Raw reads from sequencing were processed through modified DADA2 pipeline(Callahan et al. 2016) using R through an inhouse pipeline. Reads with quality score (Q score) less than 30 were removed and only high-quality reads were processed further. Filtered high quality reads were then denoised and merged using DADA2, and chimeric reads where further removed. Non-chimeric reads (i.e., amplicon sequence variant, ASVs) were then annotated using Unite (V8.2) and Silva (V138.1) databases for fungi and bacteria, respectively. All mitochondrial and chloroplast associated classifiers were eliminated and all singletons were removed from the analysis to get rid of errors due to sequencing artifacts. Altogether, there were 2.4 million reads for bacteria and 8.7 million reads for fungi. Samples were then rarefied for individual plant niches. For soil, root and leaf, each sample was rarefied at 10,000, 1,000 and 500 reads, respectively for further analysis No significant differences were observed when samples were rarefied at the same level (500 reads) for all niches.

### Statistical analysis

A flowchart representing bioinformatics and statistical approaches adopted is present in Suppl. Fig. 2. Community and biodiversity analyses were performed using the R software(R Core Team 2024). Following Mothur-based analysis, the compatible files were processed using the Phyloseq package (McMurdie and Holmes 2013). First, samples were subset into individual plant niches. Then, the alpha diversity index (Shannon index) was calculated using the VEGAN package (Oksanen et al. 2024) and differences in alpha diversities across treatments were visualised as box plots using ggplot2(Wickham 2016). ANOVA was performed to assess whether there was any significant impact of soil microbial diversity and drought treatment on microbial alpha diversity. The beta-diversity of bacterial and fungal microbial communities was assessed using Bray-Curtis distance matrices and visualised using Principal Coordinate Analysis (PCoA) plots. Global PERMANOVA analysis with 999 permutations were performed using the VEGAN package to evaluate how different variables such as plant identity (host genotype), plant niche (leaf, root, and soil), soil microbial diversity (U, 10^-2^, 10^-6^), and disturbance (drought treatment) affected the host microbiome assembly. Random Forest analysis was performed to identify key contributors of bacterial and fungal diversity from plant niche, host identity in the form of plant identity using Random Forrest package. The significance of the predicted model was then evaluated using 5000 permutations utilizing the A3 package(https://cran.r-project.org/web/packages/A3/index.html) and significance of importance of individual predictors was computed using rfPermute package (https://cran.r-project.org/web/packages/rfPermute/index.html).

Differential abundance analysis was performed using the edgeR package (Robinson et al. 2010). Samples were first subset according to plant and soil niche into leaf, root, and soil. Then, ASVs present in more than 50% of the samples were analyzed across all three soil microbial diversity gradients, and only significantly enriched ASVs (P-value >0.05) were retained for subsequent analysis. To visualise the phylogenetic relationship among these ASVs, their sequences were retrieved, aligned with MUSCLE (Edgar 2004), and phylogenetic trees were constructed using the Neighbour-Joining algorithm (Saitou and Nei 1987) using MEGA11 software (Tamura et al. 2021). The Interactive Tree of Life (iTOL) server was used to annotate the trees (Letunic and Bork 2021). Furthermore, the genera were grouped into generalist and specialist taxa for leaf microbiome to highlight the impact of host selection over leaf microbiome assembly. ASVs dominantly abundant in more than 80% of leaf samples were considered as leaf generalist and ASVs dominant for 80% of each plant species were considered as specialist taxa for that specific plant species. Finally, the Source Tracker (Knights et al. 2011) function from QIIME 1.90 (Caporaso et al. 2010) was used to build a source prediction model and to elucidate relationships between the microbiomes across different niches. This tool uses the source and sink approach to detect the possible source of microbiome based upon ASVs and metadata-based description. The soil was selected as the source for root, whereas soil and root were selected as sources for leaf microbiome. Following the prediction, models were generated for both fungi and bacteria.

### Field data survey: To establish evidence for relationship between host phylogeny and their microbiomes

To validate our glasshouse results and to ensure our findings are consistent in real world scenarios and is not impacted by soil types and climatic conditions, we obtained field data of leaf bacterial microbiomes from publicly available dataset for seven more plants (eucalypt, cotton, wheat, barley, maize, kangaroo grass, Rhodes grass and wallaby grass) where the microbiome was characterised with amplicon-sequencing using 799F/1193R primers. This dataset was a combination of data available from the European Nucleotide Archive (ENA), Sequence Read Archive (SRA) and inhouse sequencing (n=40) from other projects (Details in Supplementary Table 4). All amplicon sequencing files were processed using the same bioinformatics and statistical pipelines mentioned above to ensure that same methodology has been followed.

For all the 12 plant species (4 from glasshouse and 8 from fields), chloroplast genome sequences belonging to each of the host genera were retrieved from the NCBI genome server (https://www.ncbi.nlm.nih.gov/genome) (Supplementary Table 1). These field sites are distributed in various parts of China and Australia. Chloroplast genomic sequences were aligned using the MUSCLE algorithm (Edgar 2004) and a phylogenetic tree was generated using the Neighbour Joining algorithm with 100 bootstraps using MEGA11 (Tamura et al. 2021).To evaluate distance in their microbiome composition, hierarchical clustering was performed for microbial communities using the microbiome R package (https://bioconductor.org/packages/devel/bioc/html/microbiome.html). The topology of the plant and microbial trees was then compared to understand whether the host phylogeny and hierarchical clustering of microbiome composition had any distinct pattern across distinct species. Additionally, Spearman correlation analysis between host phylogenetic distances and their Bray-Curtis dissimilarity index was performed using the VEGAN package in R. The sequenced reads generated for this manuscript are available in NCBI (https://www.ncbi.nlm.nih.gov/sra) and can found at Sequence Read Archive (SRA) store under Bioproject PRJNA1206231.

## Results

### Host identity and niche explain plant microbiome assembly

Our result revealed that host identity and niche specification impacted the leaf and root microbiome assembly more than soil microbial diversity treatment (Figure 1). The Principal Co-ordinate analysis showed that the effect of soil initial diversity treatment on microbial community was highest for soil, followed by root, but it had minimal effect on leaf microbiomes for both fungi and bacteria (Fig. 2 a and b). PERMANOVA results supported this result (Suppl. Table 2). Random Forest based analysis revealed that, when all niches were considered together, plant niches (root and leaf) contributed significantly to the assembly of bacteria and fungi, while microbial diversity treatment had no significant impact on leaf microbiome of neither fungi nor bacteria. For individual niches, impact of host identity decreased from leaf to soil, while soil microbial diversity treatment followed the opposite trend for both fungi and bacteria. Similar trends were obtained from PERMANOVA analysis (Suppl. Table 4) and further confirmed by ordination (Fig. 2) showing a stronger impact of microbial diversity on the soil microbiome, while there was clear separation in microbial community structure with high dilution (10^-6^) treatment.

**Figure 1:**
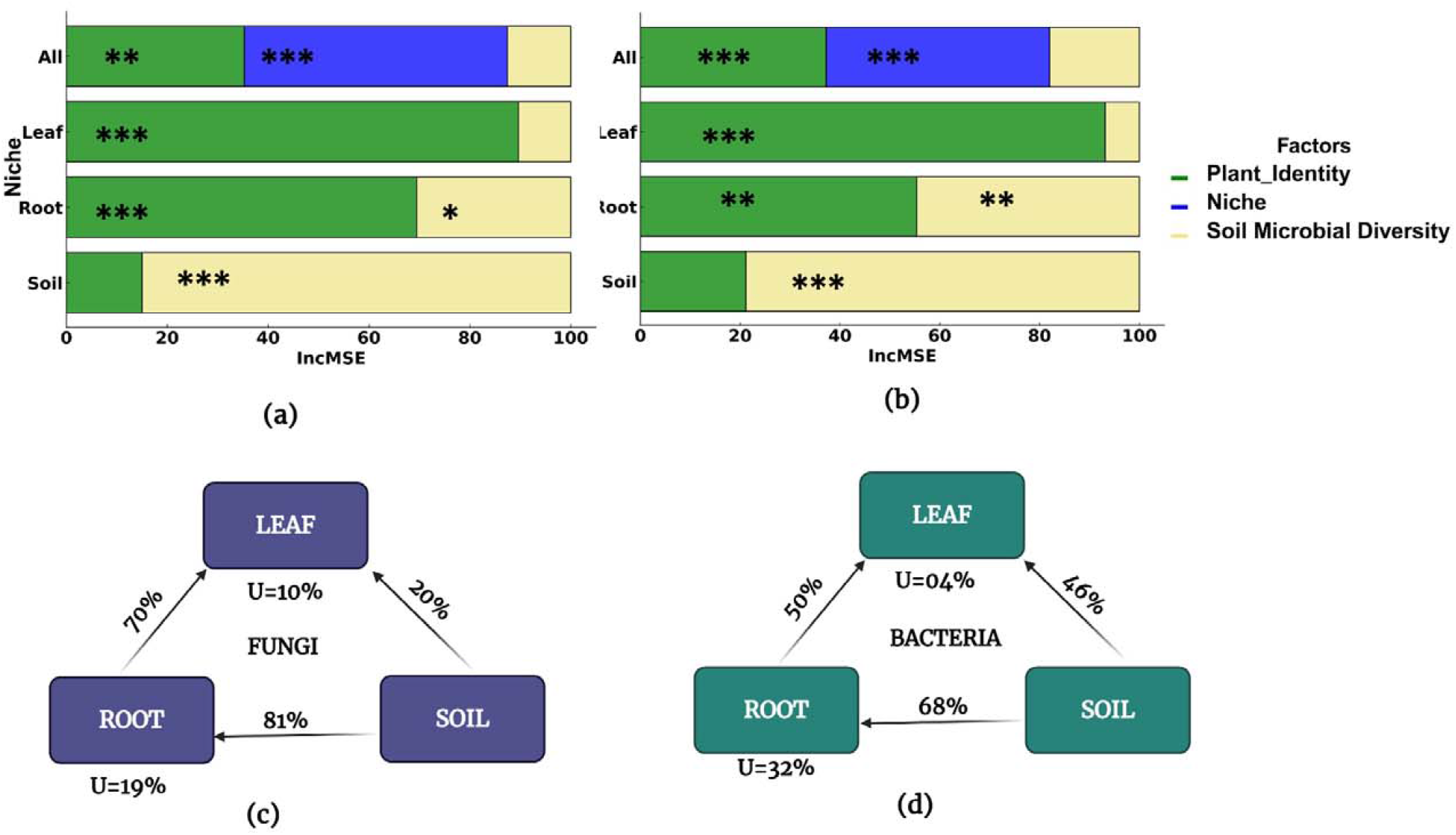
Leaf and root microbiome assemblies are mainly driven by the plant species identity (host Identity). Random Forest modelling results revealed the impact of various drivers of plant microbiome on different plant niches for (a) fungi and (b) bacteria. ‘All’ represents combined niches taken together. Plant species identity has differential impacts on microbiomes of different niches with decreasing order of Leaf >Root > Soil. Opposite trend was observed for initial soil microbial diversity treatment effects (Soil > Root > Leaf). Significance codes: \*\*\**p*□<□0.001; \*\**p*□<□0.01; \**p*□<□0.05. Plant Microbiome Source prediction models for individual plant compartments for (c) fungi and (d) bacteria. U represents an unknown source or source other than included in this study. Both source prediction models suggest that the bulk soil is main source of recruitment for the plant microbiome directly or indirectly.

**Figure 2:**
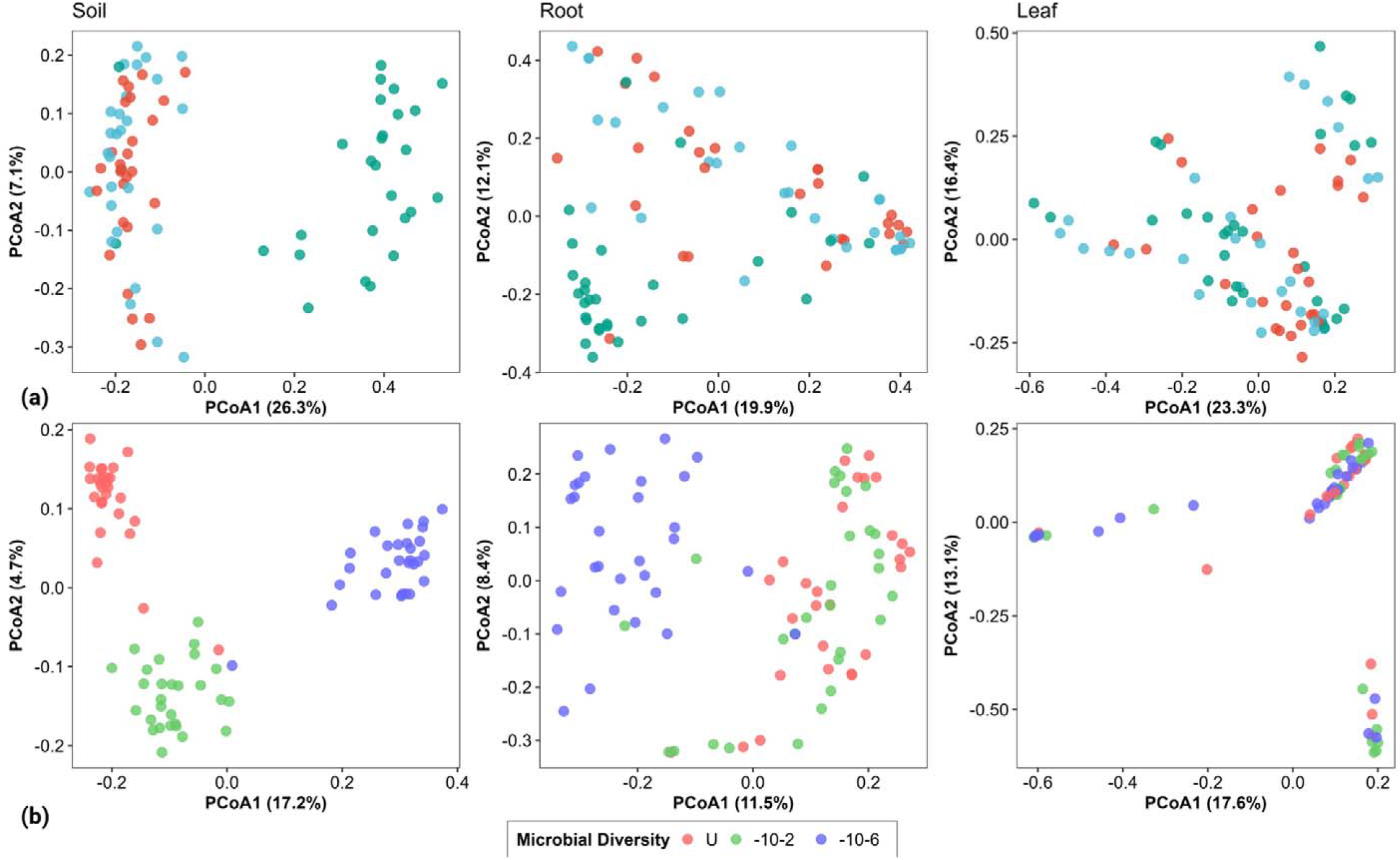
PCoA plots representing impact of soil microbial dilution treatment on microbiome assembly of different plant niches for (a) fungi and (b) bacteria. The separation between communities from leaf to soil. The host selection increased from soil to root to leaf for both bacterial and fungal communities, while initial soil microbial diversity has opposite effects. Soil diversity also created a distinct microbial community in the lowest diversity (10^-6^) soil treatment.

The source prediction model highlighted different transmission dynamics for fungi and bacteria (Fig. 1c and d). The soil fungal microbiome accounted for 81% of root endophytes and 20% of leaf fungal endophytes. About 70% of leaf fungal endophytes were potentially transmitted from the root. Our model found that root and leaf microbiomes included for 19 and 10% of unknown sources, respectively. For the root bacterial endophytes, the soil microbiome accounted for only 68% contribution as a source while the rest 32% came from the unknown source. Leaf bacterial endophytes accounted for 46% of their members from soil and 50% from the root niche.

Niche level filtering and plant identity were dominant drivers of the microbial composition. For bacteria, the leaf community was dominated by Proteobacteria (36%), Actinobacteria (30%), Firmicutes (10%), and Cyanobacteria (8%). The root community was dominated by Actinobacteria (55%), followed by Proteobacteria (35%), and Chloroflexi (2%). The soil bacterial community was mainly characterized by Actinobacteria (44%), Proteobacteria (28%), Gemmatimonadetes (4%), and Firmicutes (4%) (Suppl. Fig. 3). Ascomycota (97%) and Basidiomycota (2%) dominated the leaf fungal microbiome. Roots had a similar trend in terms of dominance, but the relative abundance of the different phyla varied, (Ascomycota = 98% and Basidiomycota = 1.2%). In soil, Ascomycota accounted for nearly 89% of the fungal sequences, followed by Mucoromycota (6%) and Mortierellomycota (2 %) (Supplementary Figure 4).

Given leaf microbiomes were distinct and less impacted by soil and environmental conditions, we examined if all taxa are unique (specialised) for individual plant species or are there taxa which are present in all/ most (generalists) of plant species. We found some taxa (Fungi: *Acremonium, Fusarium, Alternaria*, Bacteria: *Streptomyces, Bacillus, Pseudomonas, Sphingomonas, Microbacterium)* (Suppl. Table 3,4) were generalists and found in leaves of all plants and while others (Fungi: *Bisifusarium, Gliomastix, Exophilia, Exserohilum*, Bacteria: *Propionibacterineae, Angustibacter, Burkholderia, Rothia, Dyella, Reyranella, Intrasporangium, Gaiella)* were specialised taxa present only in individual plant species (Supplementary Table 3 and 4).

As expected, we found that the alpha diversity was highest in the soil, followed by the root, while the leaf harbored the lowest diversity for all plant species, supporting progressive filtering of by plants for root and leaf microbiome assemblies. The microbial diversity treatment induced soil microbial diversity gradients but impacts of initial soil microbial diversity on root microbiomes was significant for bacterial (P < 0.002) and minimal for fungal community (Suppl. Fig. 5 and 6). However, the alpha diversity of both leaf bacterial and fungal diversity was not affected by initial soil microbial diversity. There was no significant effect of drought treatment on the alpha diversity of both fungi and bacteria in any of the niches (Supplementary Figure 7 and, 8)

### Differential abundance analysis for selection of microbes by plant and soil contexts in response to initial microbial diversity gradient

The leaf-associated community had the lowest number of differentially abundant ASVs across soil microbial diversity (both bacteria (dBASVs) and fungi (dFASVs)). The taxonomic affiliation of dBASVs and dFASVs suggested a varied trend across niches. The niche-based taxonomic consistency was more evident for dBASVs. Most ASVs of a specific plant niche belonged to the same clades (Fig. 3). On the contrary, the leaf dFASVs were dispersed throughout the phylogenetic tree. The taxonomic categorization of dBASVs (Suppl. Table 3) and dFASVs supported these results (Supplementary Table 5 and 6). The common dBASVs (dBASVs 1-3) between root and soil belonged to genera *Neobacillus, Bacillus*, and *Dinococcus* respectively. Many dBASVs could not be classified taxonomically. The main representative dBASVs from leaf included *Humibacter, Xanthomonas*, and *Staphylococcus*, whereas species from *Streptomyces, Bacillus, Rhizobium*, and *Bradyrhizobium* were dominant representatives of roots. dBASVs from soil had *Bacillus* and *Rhizobium* as the most frequent genera, including specialized Rhizobium-related genera such as *Bradyrhizobium* and *Mesorhizobium*. In terms of fungi, leaf dFASVs were mainly represented by genera from *Fusarium, Fusicola*, and *Gongronella*. Shared dFASVs between all niches belonged to the genera *Humicola, Fusarium, and Purpureocillium. Fusarium* was one of the most abundant genera among dFASVs and represented multiple ASVs in all niches. Shared ASVs across root and soil were represented by numerous ascomycetous genera, including *Fusarium, Penicillium, Trichoderma, and Geomyces*

**Figure 3:**
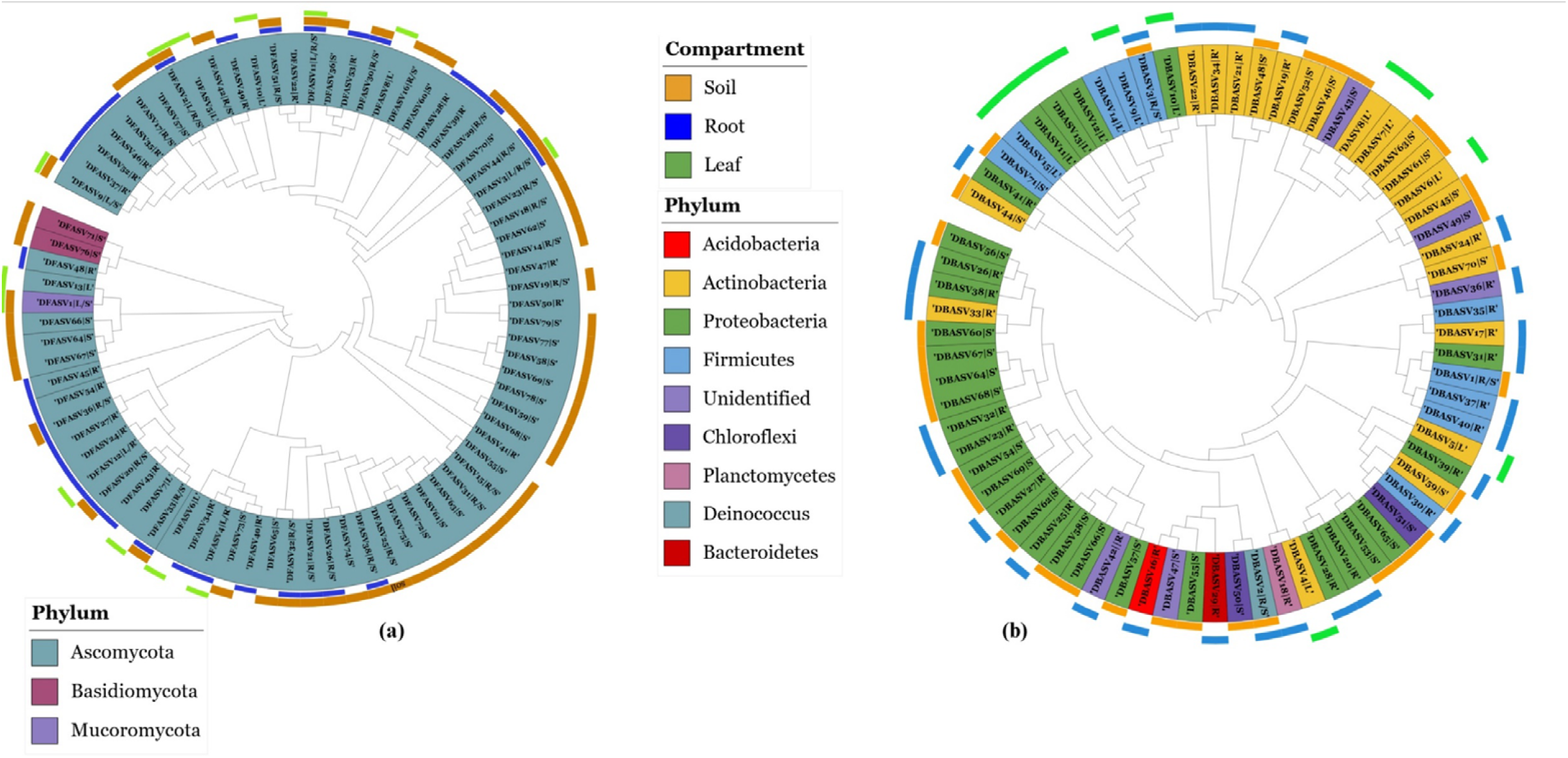
Identity of differentially abundant ASVs which were shared between different niches. Sequenced based phylogenetic tree for (a) fungi. dFASVs for leaf are phylogenetically dispersed despite belonging to the same niche. (b) for bacteria, dBASVs were, the niche-specific clades are more readily dispersed compared to fungi. This indicates that host selection has greater impact on leaf bacterial colonization compared to the fungi. Here, the phylogenetic consistency in differentially abundant ASVs suggest that bacteria with similar phylogenetic affiliation were selected by the plant despite of reduced initial pool of soil microbiome highlight dominance of host selection for the leaf microbiome.

### Leaf microbiome demonstrates linkage host phylogenetic signature

To establish whether an association exists between host phylogeny and their microbiomes, the correlation between host phylogenetic distances and their Bray-Curtis dissimilarities of leaf microbiomes were examined (Supplementary Table 7, 8). A strong positive and significant correlation was observed for both datasets, glass house experiment and field dataset (Fig. 4b,c,e). For PCoA plots of leaf fungal community, a distinct cluster for the nitrogen-fixing plant lucerne was observed and the clusters for lucerne were less distinct for root (Suppl. Fig.9 a and b). A similar trend was observed for bacteria, but the clustering was distinct for both leaf and root (Suppl. Fig. 10 a and b). Microbiome hierarchical clustering and host phylogenetic analysis revealed similar results, and plant species belonging to the same functional group was found in the same clade (Figure 4a). Here, lucerne (nitrogen-fixing) was distant from the other four plants of the C4 and C3 functional groups. The overlaying of phylogenetic distance data and microbiome data using hierarchical clustering approach showed that leaf and root bacterial communities had very distinct clusters for lucerne, and a strong and positive correlation existed between phylogenetic distance and leaf microbiome hierarchal clustering for bacteria (Spearman correlation coefficient R= 0.87, P.Value =0.002) and for fungi (Spearman correlation coefficient R= 0.75, P.Value =0.018) (Figure 4b and c ). The correlation trend was confirmed using Pearson correlation as well. There was no significant correlation observed of host phylogenetic distance with root for both bacteria and fungi.

**Figure 4:**
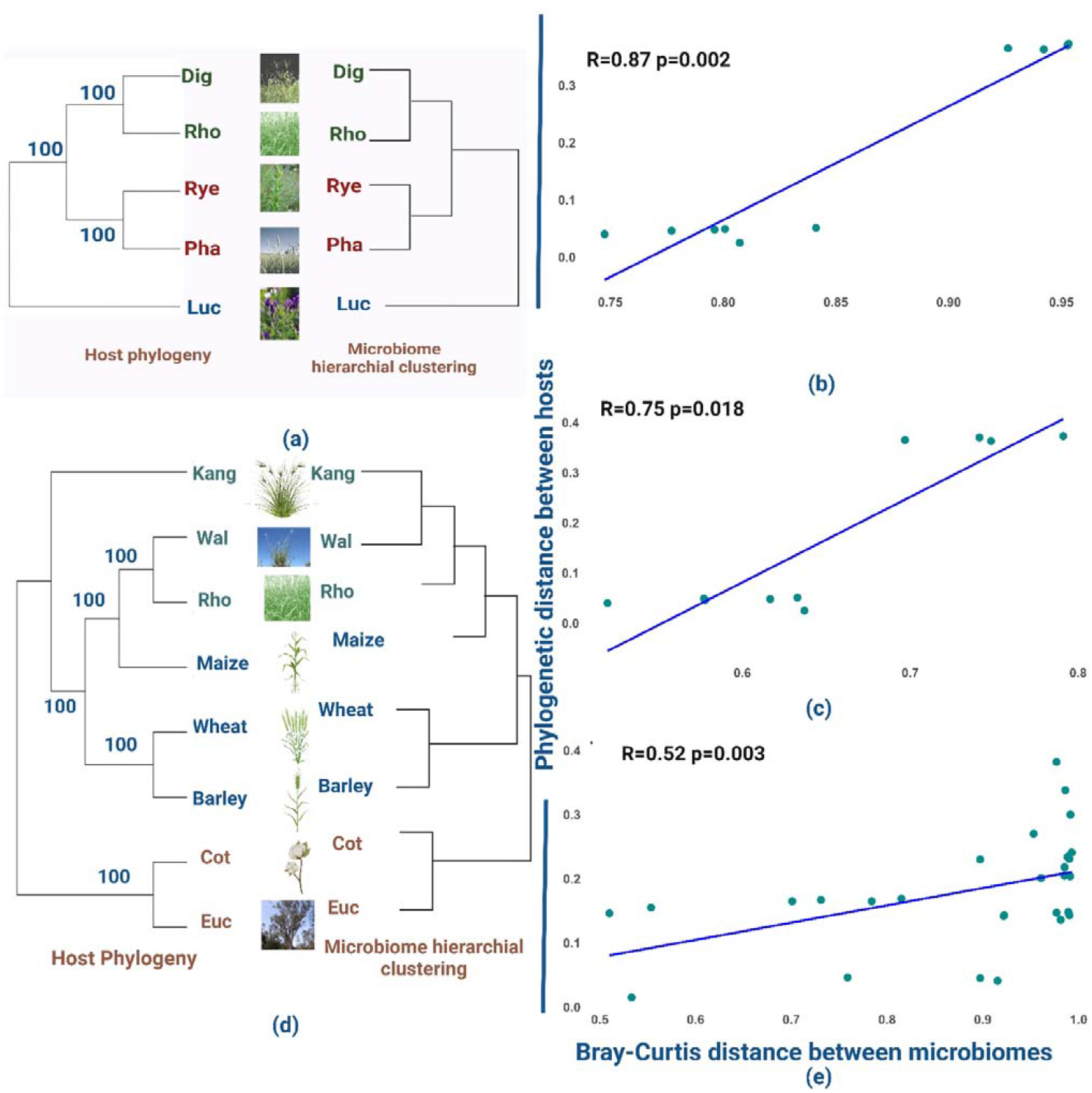
Host phylogeny and relationship between their microbiome: (a) Host phylogeny vs Microbiome hierarchal clustering dendrogram from glasshouse experiment. C3 plants and C4 plants belong to similar clades while N-fixing plant lucerne exists as an outgroup in both host phylogenetic tree. (b) Scatterplot representing correlation between Bray Curtis dissimilarity between bacterial microbiomes and corresponding phylogenetic distances of theirs host from glasshouse experiment. Spearman correlation coefficient showed strong, positive, and significant correlation. (c) Scatterplot representing correlation between Bray Curtis dissimilarity between fungal microbiomes and corresponding phylogenetic distances of theirs host from glasshouse experiment supported by Spearman correlation coefficient which showed strong, positive, and significant correlation. (d) Host phylogeny vs Microbiome hierarchal clustering dendrogram from the field survey, (e) Scatterplot representing correlation between Bray Curtis dissimilarity between microbiomes and corresponding phylogenetic distances of theirs host from additional dataset. Spearman correlation coefficient revealed strong, positive, and significant correlation between phylogenetic distance and microbiome hierarchal clustering. (Cot=cotton, Euc=eucalypt, Kang=kangaroo grass, Rho=Rhodes grass, Wal=wallaby grass.

Similar observation for plant phylogenetic distance and microbiome community relation were obtained from filed survey data (Spearman correlation coefficient R= 0.52, P.Value =0.003). (Figure 4e). Eucalyptus and cotton formed a clade in both microbiomes based dendrogram and host phylogenetic tree. Similarly, barley and wheat formed a clade while grasses (kangaroo grass, wallaby grass, Rhodes grass) formed another clade with maize plant. This pattern was almost identical for both microbiome dendrogram and host phylogenetic tree (Fig. 4d). The results were supported by ordination plot of microbiomes (Suppl. Fig. 11 and Suppl. Table 8).

## Discussions

This study revealed a strong association between host phylogeny and their microbiomes, particularly for leaf microbiomes. The Bray-Curtis distances among the microbiomes of closely related plants were similar, while more distant hosts exhibited differences in their microbiome composition. This was consistent for both glasshouse experiment and datasets obtained from field surveys. In fact, the phylogenetic distance between host plants almost mirrored the Bray-Curtis distance in microbiomes providing novel evidence for co-evolution of between plant host and its microbiome (Fitzpatrick et al. 2018; Bergelson et al. 2019; Koskella and Bergelson 2020; Zhou et al. 2023).

### Experimental evidence for dynamic co-evolutionary relationship between hosts and their microbiome

In this study, we found that plant phylogenetic distance corelated strongly with the difference in microbial compositions of different plant species and thus provided evidence in support of plant and microbiome co-evolution. Theoretical frameworks for plant -microbial interactions are well argued, given they have co-evolved together for millennia (Heckman et al. 2001; Trivedi et al., 2020). This suggested that when a new plant species evolved, it was likely that it inherited some functional traits that were directly linked to microbiome recruitment from his closest relatives (Kolodny and Schulenburg 2020). Therefore, it is expected that phylogenetic relatedness between plant species should be reflected in terms of its microbial composition. In this study, we provide novel empirical evidence from combined glasshouse and field data that the plant phylogenetic relationships reflect in the distance between associated microbiome assembly. From the glasshouse experiment this was quite evident as lucerne, a nitrogen fixing plant, had distant clusters from C3 and C4 plant. Different functional groups have different metabolic pathways, related gene functions and immunes systems that means the signal produced by plant will differ, will be sensed by different microbial species and finally plant species specific immune systems will allow colonisation by different microbiota (Singh et al. 2025) . These ultimately affect plant microbiome composition and symbiotic or mutualistic association (Copeland and Schulze-Lefert 2020; Chaudhry et al. 2021; Trivedi et al. 2020; Xiong et al. 2021a) explaining the strong differentiation by host type in our results. Differences in plant metabolisms can also create shift in plant anatomy and biology, in turn reflects in physiology and immune responsive pathways (Walker et al. 2023). In this study, these physiological and functional differences have been reflected significantly in form phylogenetic distance between plant species, as well distance between their respective microbial communities.

The glasshouse experiment of this study was designed to quantitatively examine the primary drivers of plant microbiome assembly, focusing on soil microbiome (as the main source plant microbiome), species identity, and environmental disturbance. However, this experiment did not allow us to assess if geographical location, climatic condition, or soil physico-chemical properties can also have major impacts(Trivedi et al. 2020). To address this, we used leaf microbiome data (that showed strongest relationship with host phylogeny in glasshouse experiment), from field conditions from different plant species from distinct geographical regions and different countries (e.g. Australia, China). This filed survey analysis confirmed the strong positive correlation between phylogenetic distance between plant species and distance in microbiome composition and was consistent with findings from the glasshouse experiment. Results also confirmed that plant microbial selection is independent of geographical regions, climate conditions, management practices, soil types, initial soil microbial compositions (Hamonts et al. 2018). Convergence of microbial communities according to phylogenetic relatedness of host species support the theoretical framework of plant-microbial co-evolution.

Evolutionary origin of plant species can have significant impacts on plant physiology and metabolism. For example, C4 plants have been reported to have originated and evolved from C3 plants (Osborne and Sack 2012; Christin et al. 2013), which may explain relatedness among microbiome of C3 and C4 plants species (Naylor et al. 2017). On the other hand, nitrogen fixing plants are phylogenetically and metabolically distinct (Osborne and Sack 2012; Christin et al. 2013). Their biochemistry is vey different too given their ability to fix nitrogen (with *Rhizobium* spp) and have significantly low C:N ratio. This explains the distinct microbiomes of nitrogen fixing plant from those of C3 and C4 plants, studied here (Dovrat et al. 2020). In addition to these, the plants belonging to different functional group vary significantly in terms of their root exudation under the influence of their varied metabolism, the soil environmental conditions that shape root colonization of specific microbiota (Naylor et al. 2017). Root-exudate driven microbiota recruitments, in turn is likely transmitted to other plant niches such as leaf as majority of plant microbiome is recruited from the soil, our finding is supported by previous reports (Xiong et al. 2021a; Singh et al. 2023b)..

Surprisingly, in this study the plant microbiome was not significantly affected by drought treatment, rather shaped predominantly by the host identity. Previous studies reported contrasting results where some studies found minimal effects while other reported significant impact of drought on soil and plant microbiome (Naylor et al. 2017; Vilonen et al. 2023; Fan et al. 2023; Martins et al. 2024). Our findings can be explained by the fact that there was a small impact of drought on the soil microbiome, which is a major source of plant microbiome (Trivedi et al. 2020). This is likely due to the relatively short duration of the drought treatment, indicating a certain degree of resilience in the soil microbiome to drought disturbance. Though there are evidence that prolonged drought can alter rhizosphere and root microbiome composition (Naylor et al. 2017; Santos-Medellín et al. 2021), our results suggest that short term water stress might not affect the plant microbiome assembly.

### Host associated factors are major drivers of plant microbiome assembly

Our analysis revealed that host associated factor such as host niches and host identity are major driver of plant microbiome assembly driven by host selection process (Trivedi et al., 2020). From the random forest modelling, Source tracker and PERMANOVA analyses, we established that plant sequential filtering (from soil to root to leaf) and selection mechanism is the main driver of plants microbiome assembly. The plant microbiome is a metacommunity comprised of niche-specific microbiomes that exhibit both overlaps and exclusiveness in terms of community composition (Dastogeer et al. 2020; Trivedi et al. 2020; Xiong et al. 2021a). This study reveals that host identity is stronger driver of endophyte colonization compared to soil microbial diversity, but the relative importance of these two factors is modulated by plant niches. Our findings are supported by previous studies (Xiong et al. 2021a; Singh et al. 2023b). In particular, the initial soil microbial diversity treatment had a major impact on the soil microbiome composition, while the strength of this effect was gradually diluted from soil to root and then to leaf. Conversely, host identity had the strongest influence on the leaf microbiome, followed by the roots (Wagner et al. 2016; Copeland and Schulze-Lefert 2020; Morella et al. 2020; Gao et al. 2020; Xiong et al. 2021; Singh et al. 2023b). The negligible role of soil microbial diversity and the significant variation in the leaf microbiome composition across different species collectively indicate that host genetics and the internal environment of the leaves play essential roles in the host-mediated selection process of endophytes (Liu et al. 2017; González□Teuber et al. 2021).

Source tracker analysis provided further explanation for differential microbiome assemblies in various niches. The model highlighted that soil served as the main microbial bank for plant microbiome assembly, with the root being the major direct contributor to both bacteria and fungi in leaf. As majority of microbial transmission happened from soil to root to leaf, the effect of host selection appeared to become stronger along this transmission route as many microbes are transmitted from root to leaf via plant vascular systems (Santoyo 2022).The increasing impact of host filtration or selection pressure is supported by previous reports for other plant species (Xiong et al. 2021a; Singh et al. 2023b). These transmission dynamics also explain why the host selection dominates the selection of leaf microbiota, and that this effect becomes more prominent as microbes move from soil to root and ultimately to the leaf (Xiong, et al. 2021a; Xiong, et al. 2021b; Singh et al. 2023b).

Despite this strong plant identity effect, several microbiome members were consistently present in leaf microbiome across the soil dilution treatments and plant species. For example, in case of fungi, the differential abundance analysis shows that ASVs belonging to species from *Fusarium, Fusicola, Gongronella, Pyrenophora, Epicoccum, Exophiala* genera were enriched in leaf across all soil microbial diversity gradients and plant hosts, supporting the idea of generalists’ leaf taxa that occupies multiple hosts that evolved to colonise many plant species. In case of bacteria, enriched generalist in leaves were *Humibacter, Xanthomonas, Cetalospora* and *Staphylococcus*, supporting that these taxa were actively recruited through soil to leaf despite initial differences in the plant identity and initial soil microbiome source. However, our study indicates that the filtering mechanisms also facilitate colonization by selected specialist taxa.. During filtration procedure, plant species specific selection could remove incompatible microbial taxa via immune response, or those microbial taxa cannot colonise due to lack of ability to respond to chemical stimuli, ability to compete with other microbiota (Trivedi et al. 2020). There were several specialist taxa for individual plants species such as *Kocuria, Gardenella, Bisifusarium* (Lucerne), *Pseudolabrys, Strentrophomonas*, (Digitaria), *Gliomastix, Angustibacter* (Ryegrass) suggesting importance of host selection and filtering mechanisms driven by host associated features (Xiong, et al. 2021a; Xiong, et al. 2021b; Singh et al. 2023b) for selection of specialist microbiota. Overall, this support the notion that a “core” microbiome exists within plants (Hamonts et al. 2018; Toju et al. 2018; Trivedi et al. 2020) but these include both generalists and specialists taxa.

## Conclusion

Overall, our results demonstrate that leaf endophyte and root endophyte assemblies are largely influenced by host identity, host selection and host-mediated filtration mechanisms. We also provide evidence that despite being major source of indirect or direct recruitment for microbiome, soil microbial diversity and composition had no significant effect on leaf microbiome assembly. Importantly, these results provide novel evidence of broad-scale relationship between host phylogeny and variation in leaf endophytic microbiomes providing empirical evidence in support of plant-microbiome co-evolutions. These findings significantly advance our understanding on drivers of plant microbiome assembly and provide new knowledge that can be harnessed in future for improving plant productivity.

## Acknowledgements

This experiment was supported by the Australian Research Council (DP230101448). P.K.S. acknowledges support from Western Sydney University for post graduate research scholarship. EE is supported by the Australian Research Council DECRA Fellowship (DE210101822).

## Authors contributions

BKS, PKS, CSCM, EE, RHJ, PBR and CAM conceived the design the experiment. Experiment was established and maintained by CSCM and RHJ. Lab work and most of bioinformatics and statistical analyses was carried by PKS. BKS, MD-B, EE, JW also help with statistical analyses. BB, CX, GQ provided additional data. PKS written the first draft of the manuscript in close consultation with BKS and all authors edited and commented to improve the manuscript.

## Notes

### Competing Interest Statement

The authors have declared no competing interest.

