## Supplementary figures and tables for "Leaf microbiome assembly is linked to plant phylogeny"

Supplementary files:

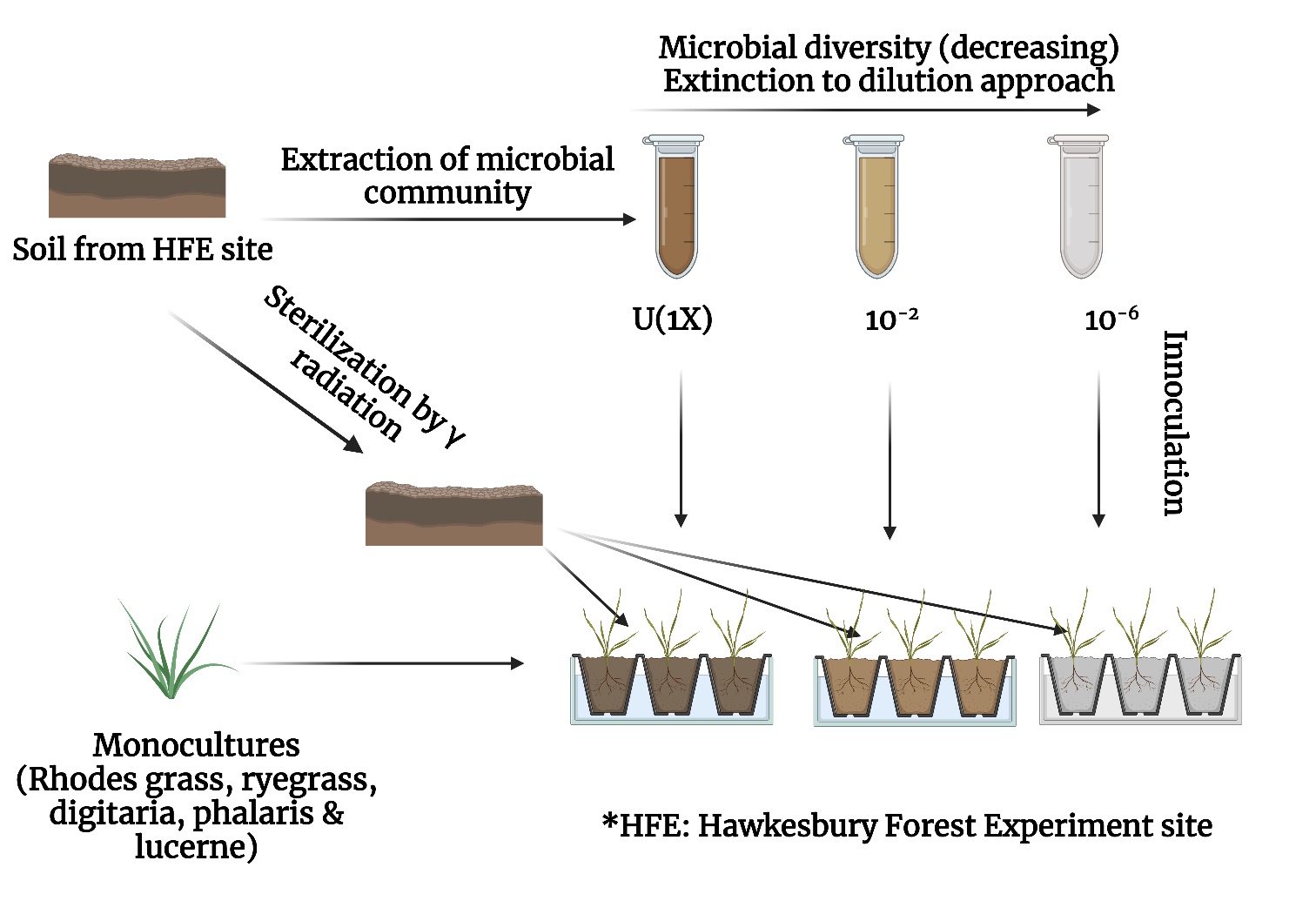

**Supplementary Figure 1**: schematic representation of the experimental designed established to test the effect of initial soil microbial diversity treatment and host identity on plant microbiome assembly of five different plant species.

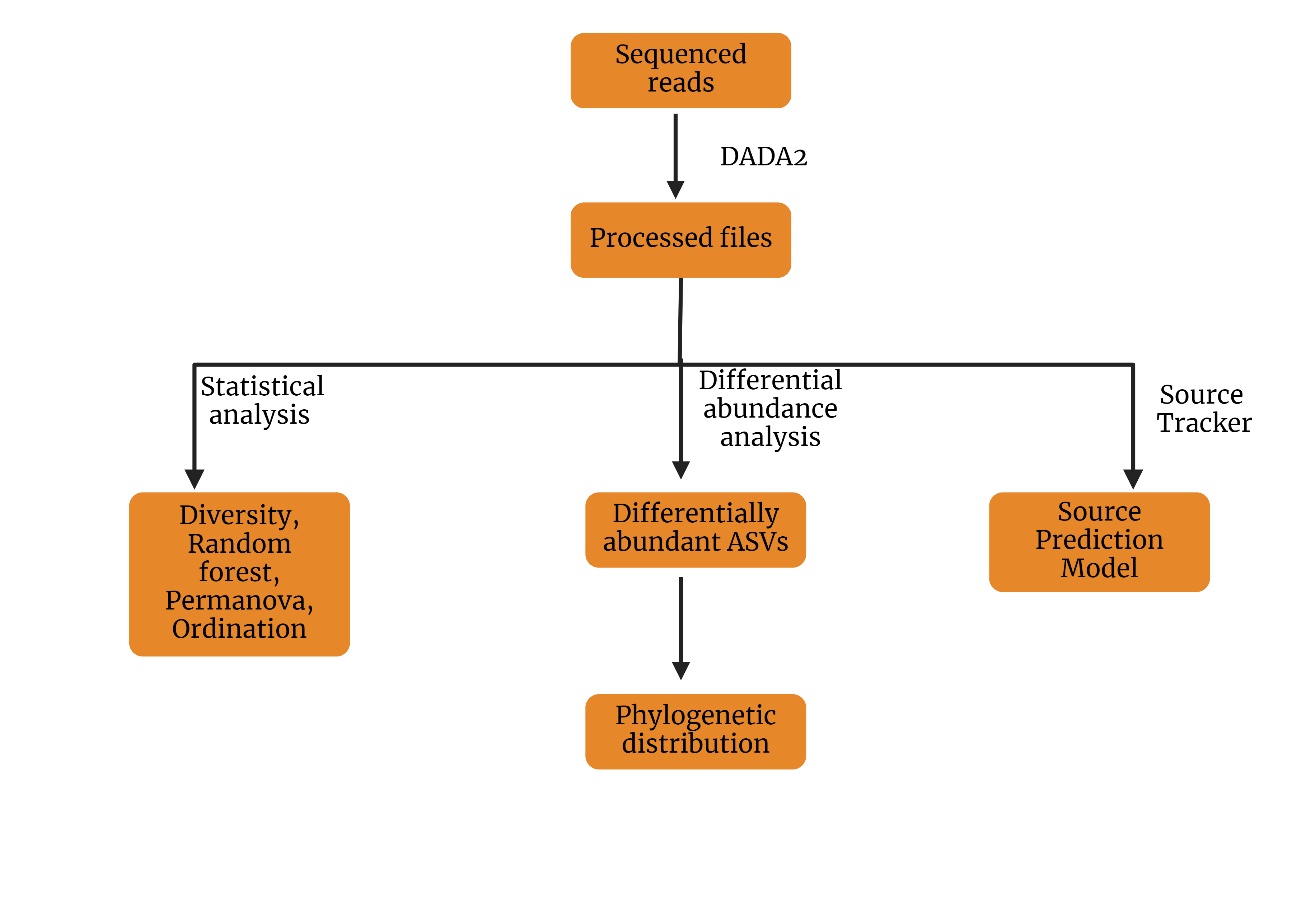

**Supplementary Figure 2:** A flowchart representing various statistical and computational approaches adopted for data analysis.

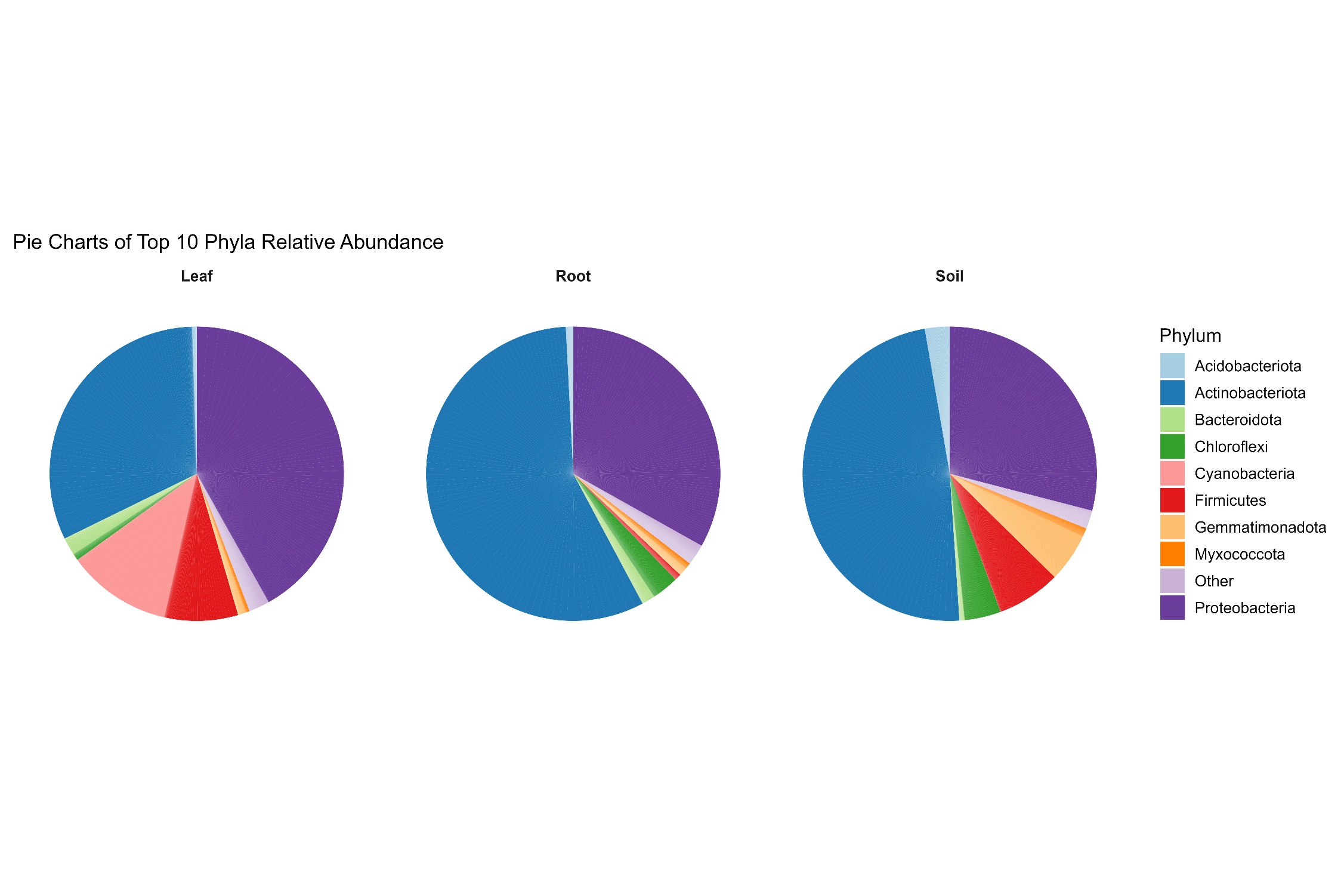

**Supplementary Figure 3**: Pie chart representing phylum level plant compartment-specific composition for bacteria (a) Leaf, (b) Root, and (c) Soil. There was variation in terms of relative abundance of different phyla across these compartments.

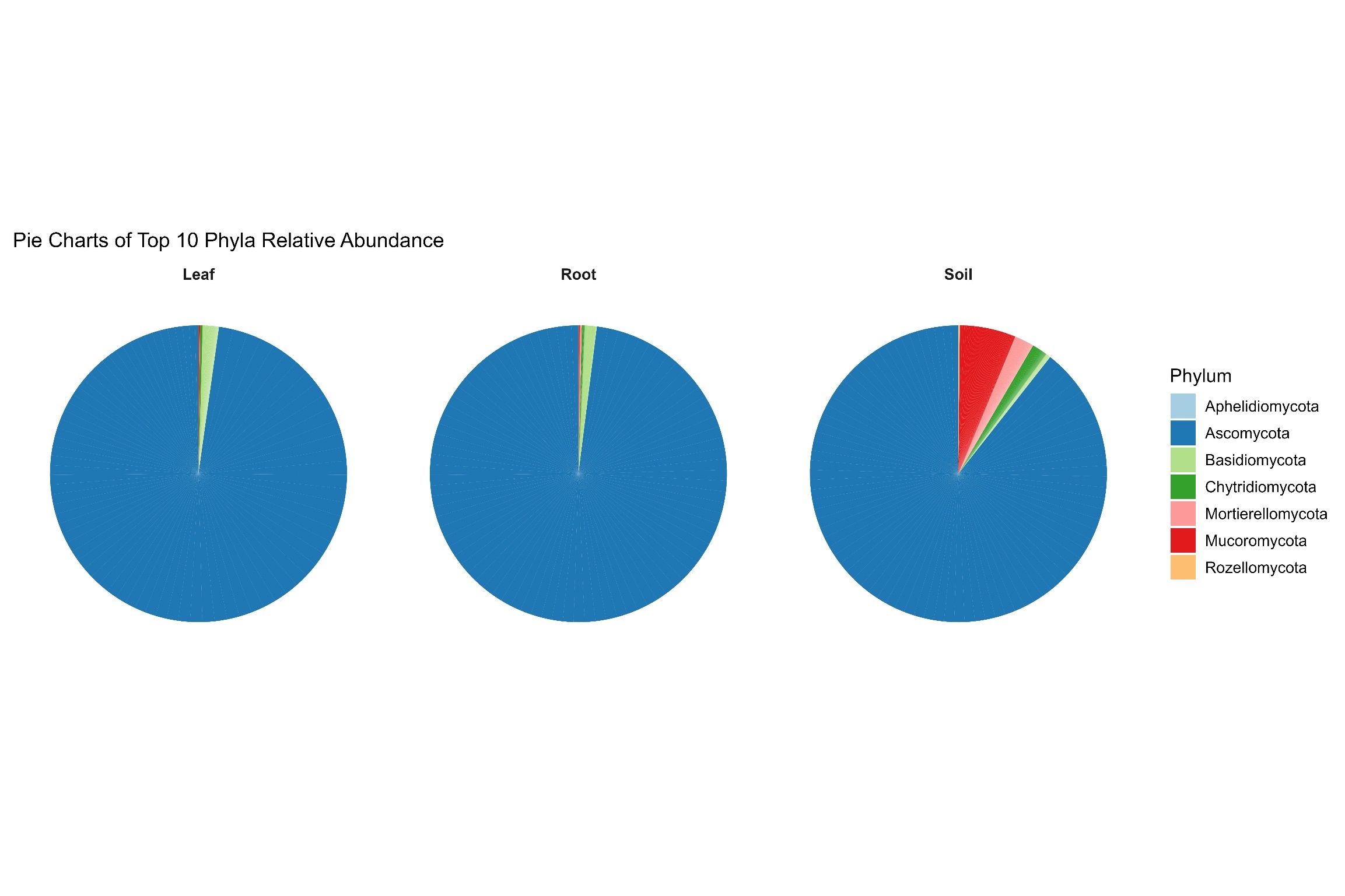

**Supplementary Figure 4**: Pie-chart representing phylum level plant compartment-specific composition for fungi (a) Leaf, (b) Root, and, (c) Soil. There was variation in terms of relative abundance of different phyla across these compartments.

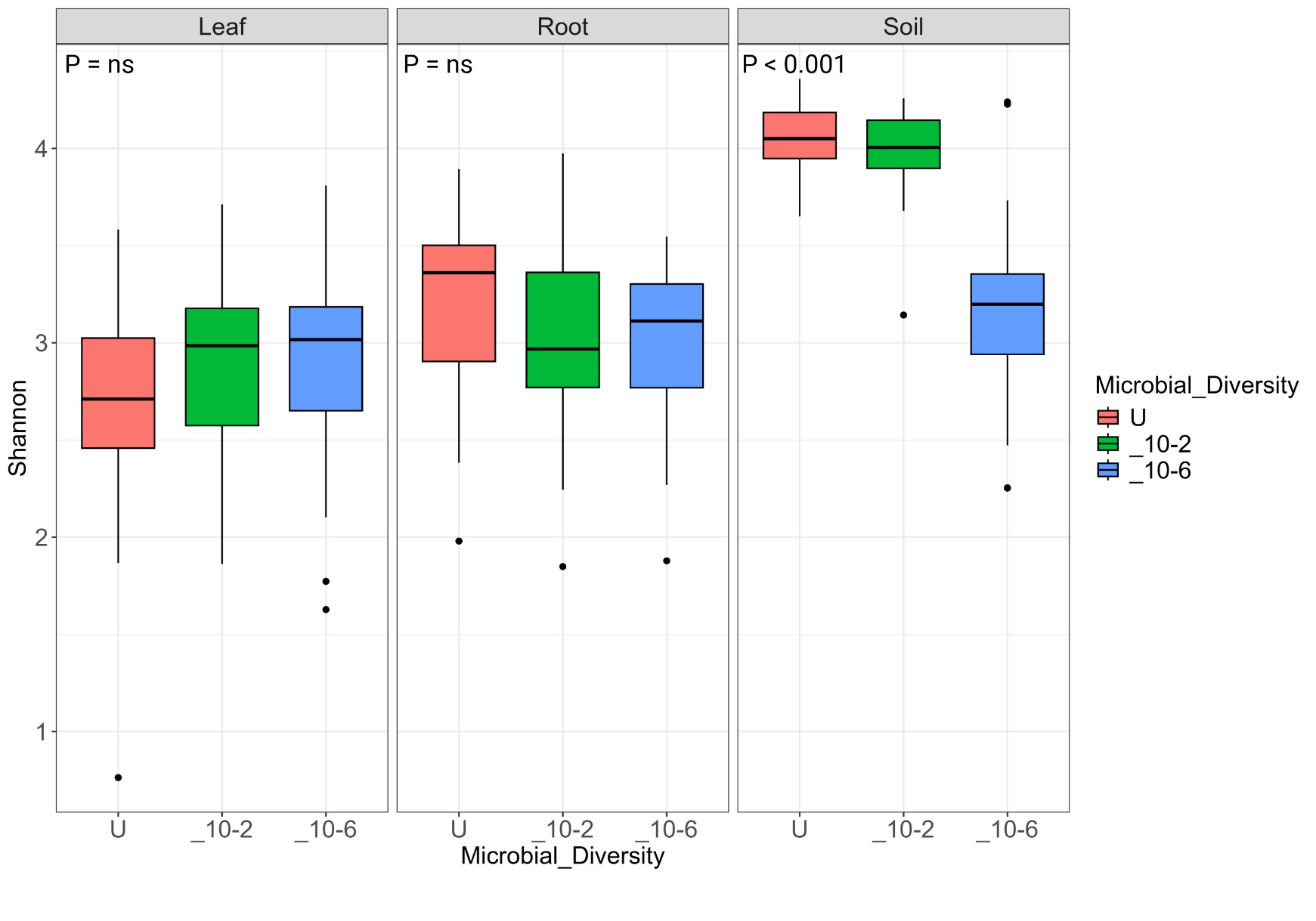

**Supplementary Figure 5**: Box plot representing the effect of soil microbial diversity over alpha diversity of fungi of different plants niches. Microbial diversity treatment significantly impacts the soil while there is no significant impact on root and leaf Shannon index. In terms overall diversity soil has highest diversity while leaf has the lowest.

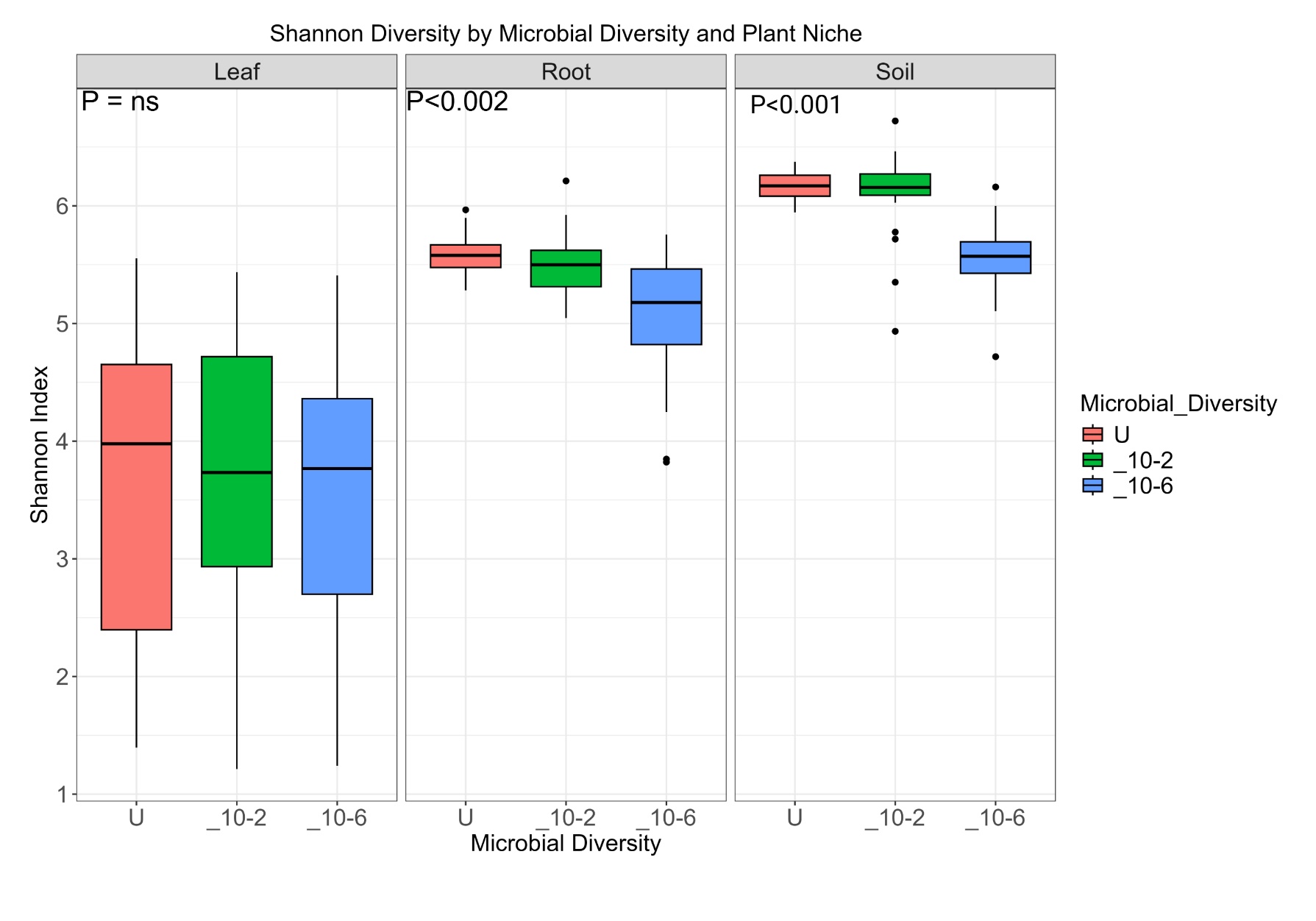

**Supplementary Figure 6**: Box plot representing the effect of soil microbial diversity over alpha diversity of bacteria of different plants and their respective plant compartments. Soil and root are significantly affected niche whereas there is no significant impact on the leaf microbiome. In terms overall diversity soil has highest diversity while leaf has the lowest.

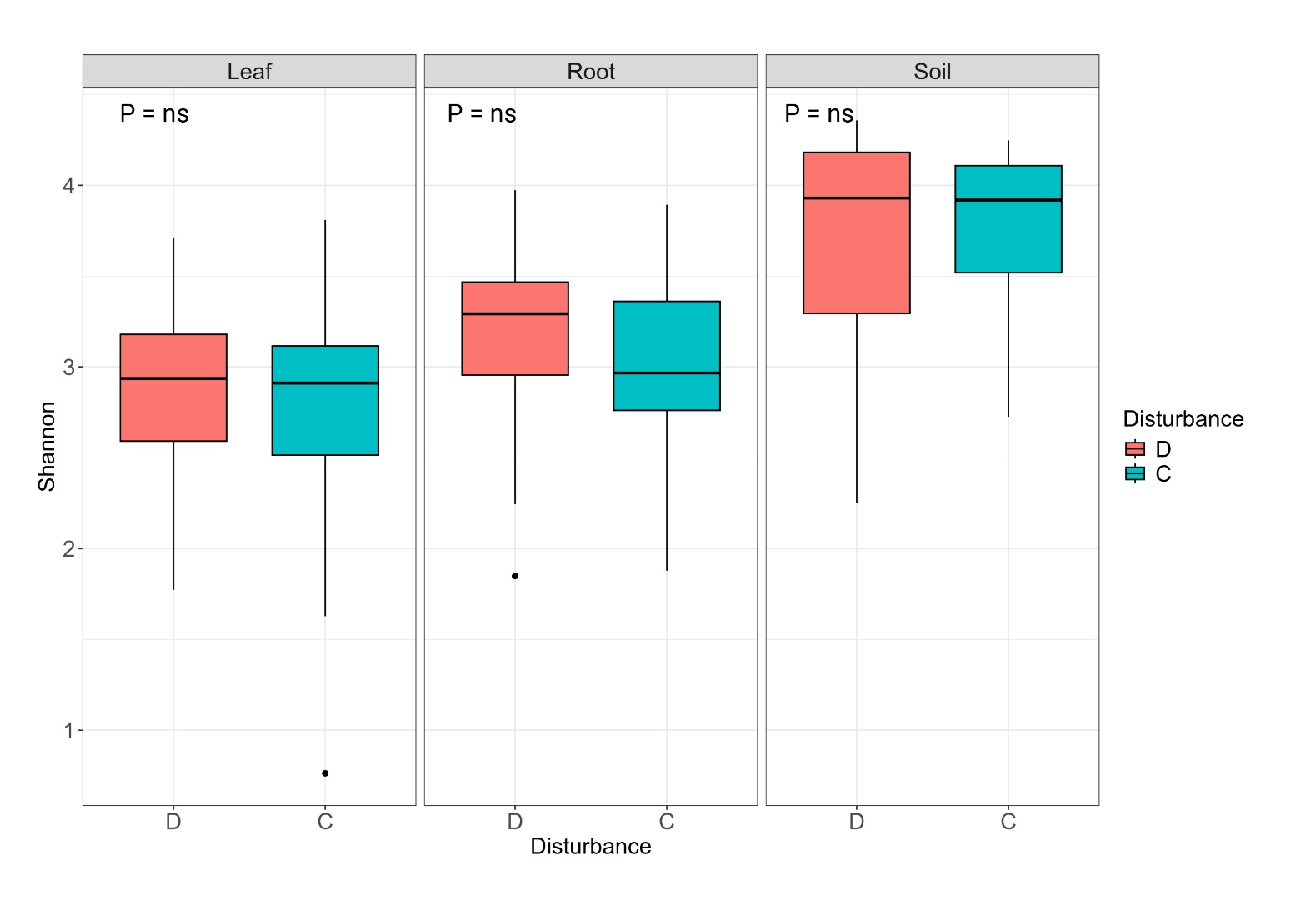

**Supplementary Figure 7**: Box plot representing the effect of disturbance over alpha diversity of fungi of different plants niches. There is no significant impact of disturbance on any of the plant niches.

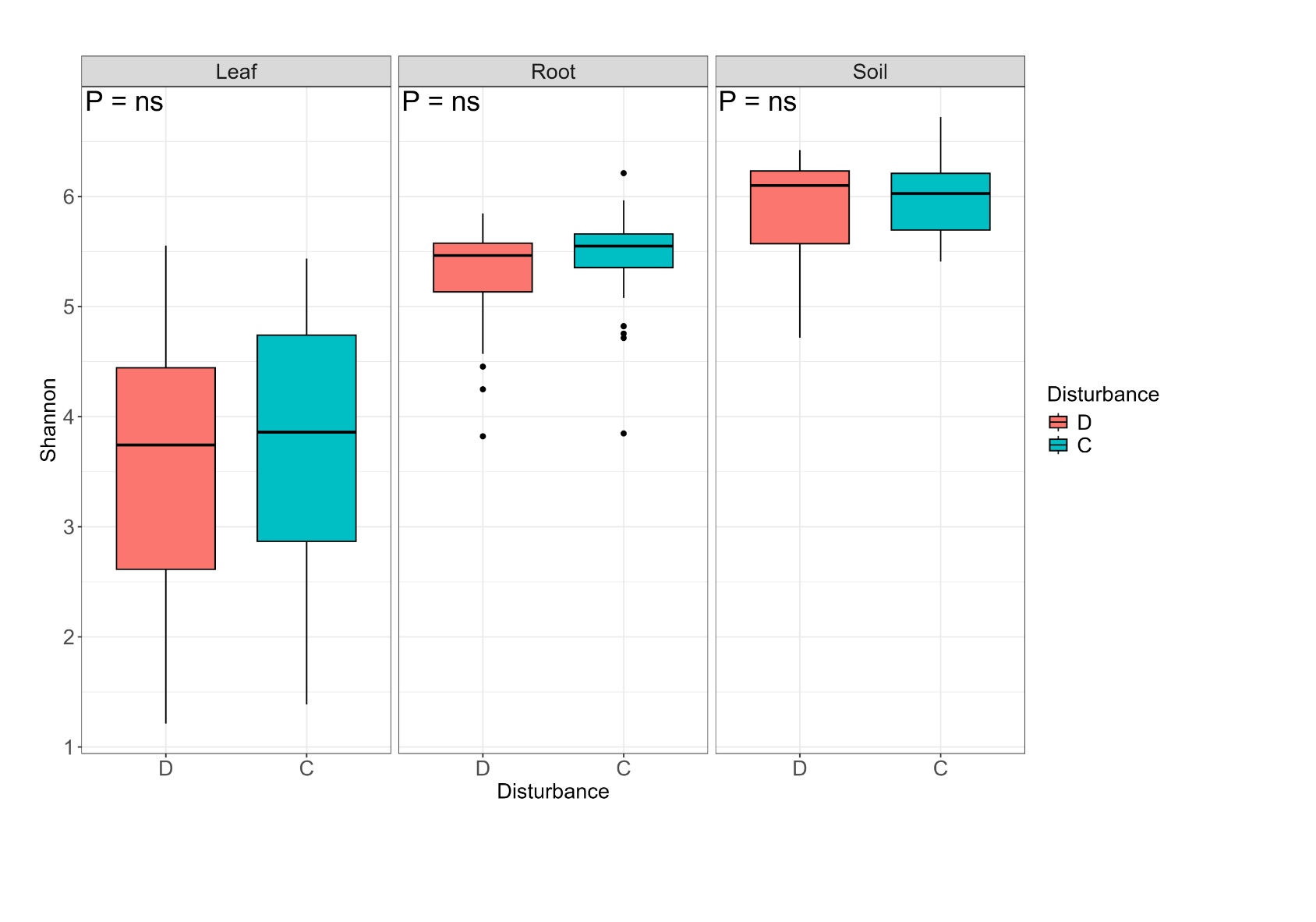

**Supplementary Figure 8**: Boxplot representing the effect of drought treatment over the alpha diversity of bacteria microbiome. There is a clear indication that disturbance in the form of drought has not significantly affected the alpha diversity.

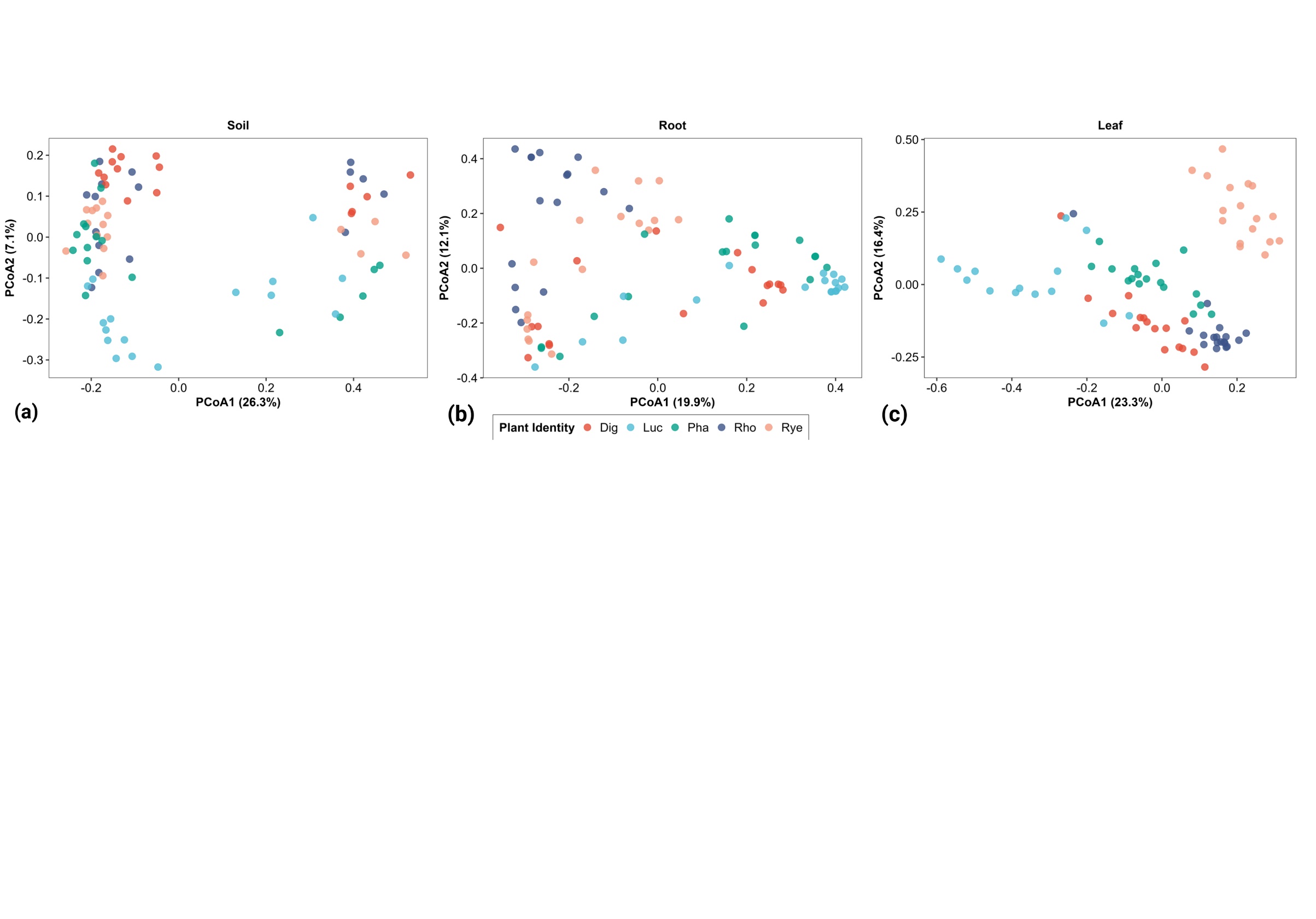

**Supplementary Figure 9**: PCoA plot representing effect of plant identity in fungi in different niche (a) soil, (b) root, and (c) leaf. Lucerne a nitrogen fixing plant forms a separate cluster from other plants in leaf and root, but it less distinct in soil.

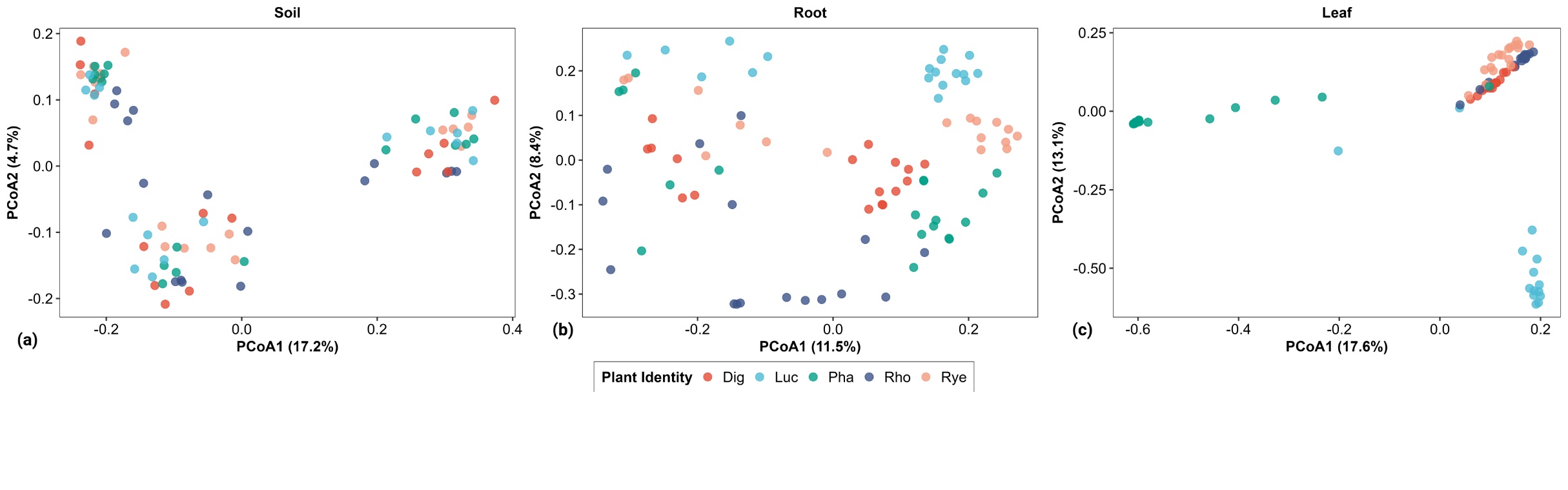

**Supplementary Figure 10:** PCoA plot representing effect of plant identity over bacteria in different niche (a) leaf, (b) root, and (c) soil. Lucerne a nitrogen fixing plant forms a separate cluster from other plants in leaf and root, but it less distinct in soil

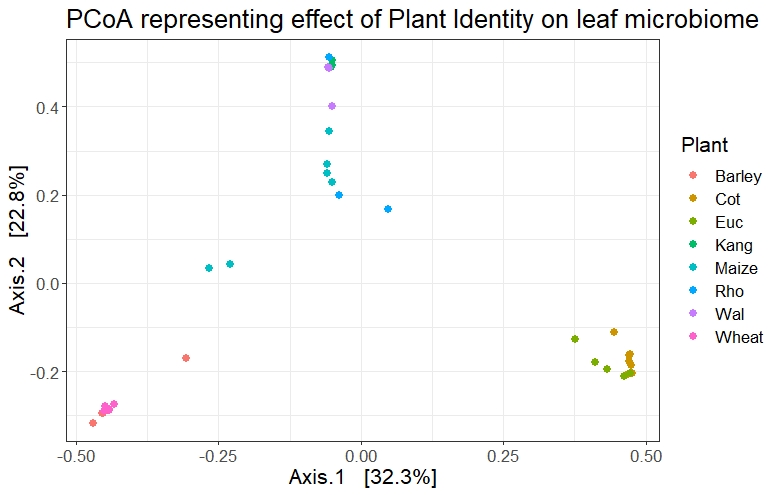

**Supplementary Figure 11**: PCoA plot representing PCoA plot representing effect of plant identity over bacterial leaf microbiome from additional dataset analysed. (Cot=cotton, Euc=Eucalypt, Kang=kangaroo grass, Rho=Rhodes grass, Wal=wallaby grass,)

Supplementary Tables

**Supplementary Table 1**: Accession numbers of chloroplast genomes and additional amplicon sequencing datasets used in the experiment.

| Serial No | Plant | NCBI accession |
| --- | --- | --- |
| 1 | Phalaris cpgenome | NC_027481.1 |
| 2 | Digitaria cpgenome | NC_024176.1 |
| 3 | Ryegrass cpgenome | NC_009950.1 |
| 4 | Rhodes grass cpgenome | NC_042841.1 |
| 5 | Lucerene cpgenome | NC_042841.1 |
| 6 | Wheat cpgenome | NC_002762.1 |
| 7 | Eucalypt cpgenome | NC_022397.1 |
| 8 | Maize cpgenome | NC_001666.2 |
| 9 | Barley cpgenome | NC_008590.1 |
| 10 | Kangaroo grass cpgenome | NC_035016.1 |
| 11 | Wallaby grass cpgenome | NC_037166.1 |
| 12 | Cotton cpgenome | NC_007944.1 |
| Amplicon sequencing data | | |
| 1 | Wheat, Barley, Maize (Origin : China) | SRA: PRJNA559707 |
| 2 | Eucalypt (Origin : Australia) | In house |
| 3 | Cotton (Origin : Australia, multiple sampling sites) | In house |
| 4 | Kangaroo grass. Wallaby grass, Rhodes grass (Origin : Australia) | ENA: PRJEB38041 |

**Supplementary Table 2:** Table representing R^2^ values from Global PERMANOVA analysis for the impact of different plant compartments, plant combinations, soil microbial diversity, and disturbance over bacterial and fungal diversity.

|  |  | **Fungi** | **Bacteria** |
| --- | --- | --- | --- |
| **Features** | **Factors** | **R2** | **R2** |
| **Ryegrass** | Niche | 0.459941 | 0.161712 |
|  | Disturbance | 0.007545 | 0.016711 |
|  | Soil Microbial Diversity | 0.09586 | 0.058769 |
| **Rhodes** | Niche | 0.476394 | 0.433139 |
|  | Disturbance | 0.004496 | 0.008615 |
|  | Soil Microbial Diversity | 0.112705 | 0.057369 |
| **Phalaris** | Niche | 0.542653 | 0.424078 |
|  | Disturbance | 0.006434 | 0.014013 |
|  | Soil Microbial Diversity | 0.078307 | 0.059703 |
| **Digitaria** | Niche | 0.441929 | 0.580348 |
|  | Disturbance | 0.004311 | 0.010208 |
|  | Soil Microbial Diversity | 0.066737 | 0.046192 |
| **Lucerne** | Niche | 0.45253 | 0.389396 |
|  | Disturbance | 0.008948 | 0.014013 |
|  | Soil Microbial Diversity | 0.07006 | 0.048051 |
| **Soil** | Plant Identity | 0.104973 | 0.062379 |
|  | Disturbance | 0.005385 | 0.009785 |
|  | Soil Microbial Diversity | 0.289991 | 0.28963 |
| **Root** | Plant Identity | 0.202797 | 0.192307 |
|  | Disturbance | 0.006406 | 0.008187 |
|  | Soil Microbial Diversity | 0.075218 | 0.159486 |
| **Leaf** | Plant Identity | 0.258352 | 0.226308 |
|  | Disturbance | 0.009062 | 0.010509 |
|  | Soil Microbial Diversity | 0.027808 | 0.039678 |
| **All** | Plant Identity | 0.103914 | 0.088831 |
|  | Niche | 0.158316 | 0.280801 |
|  | Disturbance | 0.001694 | 0.001532 |
|  | Soil Microbial Diversity | 0.029123 | 0.021856 |

**Supplementary Table 3**: Generalist and specialist taxa fungi.

| **Generalist** | **Specialist taxa Digitaria** | **Specialist taxa Lucerne** | **Specialist taxa Rhodes** | **Specialist taxa Ryegrass** | **Specialist taxa Phalaris** |
| --- | --- | --- | --- | --- | --- |
| *Acremonium* | *Cladosporium* | *Cladosporium* | *Cladosporium* | *Cladosporium* | *Cladosporium* |
| *Alternaria* | *Fusarium* | *Fusarium* | *Fusarium* | *Fusarium* | *Fusarium* |
| *Chaetomium* | *Paraconiothyrium* | *Paraconiothyrium* | *Paraconiothyrium* | *Paraconiothyrium* | *Paraconiothyrium* |
| *Cladosporium* | *Myrmecridium* | *Myrmecridium* | *Myrmecridium* | *Myrmecridium* | *Myrmecridium* |
| *Clonostachys* | *Edenia* | *Purpureocillium* | *Edenia* | *Edenia* | *Edenia* |
| *Coniochaeta* | *Poaceascoma* | *Humicola* | *Poaceascoma* | *Poaceascoma* | *Poaceascoma* |
| *Dictyochaeta* | *Pseudorhypophila* | *Pseudothielavia* | *Pseudorhypophila* | *Pseudorhypophila* | *Pseudorhypophila* |
| *Edenia* | *Purpureocillium* | *Fungi_gen_Incertae_sedis* | *Purpureocillium* | *Purpureocillium* | *Purpureocillium* |
| *Epicoccum* | *Humicola* | *Edenia* | *Humicola* | *Humicola* | *Humicola* |
| *Fungi_gen_Incertae_sedis* | *Pseudothielavia* | *Pseudorhypophila* | *Pseudothielavia* | *Pseudothielavia* | *Pseudothielavia* |
| *Fusarium* | *Talaromyces* | *Verticillium* | *Verticillium* | *Talaromyces* | *Talaromyces* |
| *Gongronella* | *Verticillium* | *Epicoccum* | *Epicoccum* | *Verticillium* | *Epicoccum* |
| *Humicola* | *Epicoccum* | *Alternaria* | *Sarocladium* | *Epicoccum* | *Alternaria* |
| *Myrmecridium* | *Alternaria* | *Neocosmospora* | *Alternaria* | *Sarocladium* | *Pseudopithomyces* |
| *Neocosmospora* | *Neocosmospora* | *Penicillium* | *Neocosmospora* | *Alternaria* | *Clonostachys* |
| *Paraconiothyrium* | *Clonostachys* | *Pseudopithomyces* | *Penicillium* | *Neocosmospora* | *Acremonium* |
| *Penicillium* | *Pseudopithomyces* | *Clonostachys* | *Chaetomium* | *Clonostachys* | *Gongronella* |
| *Poaceascoma* | *Penicillium* | *Trichoderma* | *Pseudopithomyces* | *Chaetomium* | *Scytalidium* |
| *Pseudopithomyces* | *Chaetomium* | *Gongronella* | *Clonostachys* | *Pseudopithomyces* | *Rhizoctonia* |
| *Pseudorhypophila* | *Trichoderma* | *Dictyochaeta* | *Trichoderma* | *Acremonium* | *Sarocladium* |
| *Pseudothielavia* | *Gongronella* | *Coniochaeta* | *Dictyochaeta* | *Rhizoctonia* | |
| *Purpureocillium* | *Acremonium* | *Chaetomium* | *Gongronella* | *Exophiala* | |
| *Sarocladium* | *Dictyochaeta* | *Bisifusarium* | *Coniochaeta* |  |  |
| *Talaromyces* | *Sarocladium* | *Sarocladium* | *Gliomastix* |  |  |
| *Trichoderma* | *Exophiala* | *Fusicolla* |  |  |  |
| *Verticillium* | *Fusicolla* | *Exserohilum* |  |  |  |
|  |  | *Exophiala* |  |  |  |

**Supplementary Table 4**: Generalist and specialist taxa of bacteria.

| **Generalist** | **Specialist taxa Ryegrass** | **Specialist taxa Rhodes** | **Specialist taxa Lucerne** | **Specialist taxa Phalaris** | **Specialist taxa Digitaria** |
| --- | --- | --- | --- | --- | --- |
| *Streptomyces* | *Micrococcineae* | *Streptomyces* | *Streptomyces* | *Streptomyces* | *Amycolatopsis* |
| *Amycolatopsis* | *Propionibacterineae* | *Amycolatopsis* | *Amycolatopsis* | *Amycolatopsis* | *Streptomyces* |
| *Curtobacterium* | *Corynebacterineae* | *Pedococcus-Phycicoccus* | *Mesorhizobium* | *Curtobacterium* | *Pedococcus-Phycicoccus* |
| *Mesorhizobium* | *Streptococcus* | *Curtobacterium* | *Bacillus* | *Mesorhizobium* | *Curtobacterium* |
| *Nocardioides* | *Pseudomonas* | *Mesorhizobium* | *Nocardiodes* | *Rhodococcus* | *Mesorhizobium* |
| *Bacillus* | *Chryseobacterium* | *Bacillus* | *Geodermatophilus* | *Microbacterium* | *Nocardioides* |
| *Geodermatophilus* | *Amycolatopsis* | *Geodermatophilus* | *Cutibacterium* | *Dyella* | *Bacillus* |
| *Actinoplanes* | *Rhodococcus* | *Actinoplanes* | *Microbacterium* | *Cutibacterium* | *Rhodococcus* |
| *Dyella* | *Actinoplanes* | *Rhodococcus* | *Dyella* | *Ralstonia* | *Actinoplanes* |
| *Rhodococcus* | *Pseudonocardia* | *Microbacterium* | *Ralstonia* | *Lysinimonas* | *Sphingomonas* |
| *Bacillus* | *Bacillus* | *Dyella* | *Kocuria* | *Deinococcus* | *Intrasporangium* |
| *Actinoplanes* | *Rhodococcus* | *Intrasporangium* | *Rothia* | *Pseudonocardia* | *Cutibacterium* |
| *Intrasporangium* | *Geodermatophilus* | *Cutibacterium* | *Gardnerella* | *Methylobacterium* | *Pseudolabrys* |
| *Mycobacterium* | *Pseudarthrobacter* | *Nocardioides* | *Spingomonas* | *Staphylococcus* | *Microbacterium* |
| *Microbacterium* | *Terrabacter* | *Ralstonia* | *Deinococcus* | *YC-ZSS-LKJ147* | *Geodermatophilus* |
| *Dyella* | *Pantoea* | *Burkholderia* | *Burkholderia* | *Williamsia* | *Pseudonocardia* |
| *Nocardioides* | *Pedococcus* | *Pseudonocardia* | *Reyranella* | *Pseudomonas* | *Reyranella* |
| *Ralstonia* | *Microbacterium* | *Terrabacter* | *Rothia* | *Sphingomonas* | *Terrabacter* |
| *Pseudonocardia* | *Arthrobacter* | *Arthrobacter* | *Reyranella* | *Aeromicrobium* | *Pseudarthrobacter* |
| *Bacillus* | *Angustibacter* | *Dokdonella* |  | *Spirosoma* | *Dyella* |
| *Dokdonella* | *Cutibacterium* | *Pseudarthrobacter* |  | *Marmoricola* | *Stenotrophomonas* |
| *Geodermatophilus* | | *Pantoea* |  |  | *Staphylococcus* |
| *Sphingomonas* |  | *Altererythrobacter* |  |  | *Corynebacterium* |
| *Reyranella* |  | *Leifsonia* |  |  | *Deinococcus* |
| *Chelativorans* |  |  |  |  | *Brevundimonas* |
|  |  |  |  |  | *Paracoccus* |
|  |  |  |  |  | *Gaiella* |

**Supplementary Table 5:** Differentially abundant fungal ASVs (dFASVs), their plant compartments, phylum, and genus

| **ID** | **Phylum** | **Genus** | **Compartment** |
| --- | --- | --- | --- |
| dFASV1 | Mucoromycota | *Gongronella* | Leaf and Soil |
| dFASV2 | Ascomycota | *Fusarium* | Leaf, Root, and Soil |
| dFASV3 | Ascomycota | *Humicola* | Leaf, Root, and Soil |
| dFASV4 | Ascomycota | *Exophiala* | Leaf and Root |
| dFASV5 | Ascomycota | *Fusicola* | Leaf |
| dFASV6 | Ascomycota | *Ramichlorodium* | Leaf |
| dFASV7 | Ascomycota | *Uncultured Ascomycota* | Leaf |
| dFASV8 | Ascomycota | *Emericellopsis* | Leaf |
| dFASV9 | Ascomycota | *Pyrenophora* | Leaf and Soil |
| dFASV10 | Ascomycota | *Uncultured Ascomycota* | Leaf |
| dFASV11 | Ascomycota | *Romanomerm* | Leaf, Root, and Soil |
| dFASV12 | Ascomycota | *Epicoccum* | Leaf and Root |
| dFASV13 | Ascomycota | *Gongronella* | Leaf |
| dFASV14 | Ascomycota | *Chaetomium* | Root and Soil |
| dFASV15 | Ascomycota | *Cladosporium* | Root and Soil |
| dFASV16 | Ascomycota | *Clonostachys* | Root and Soil |
| dFASV17 | Ascomycota | *Fusarium* | Root and Soil |
| dFASV18 | Ascomycota | *Pseudothielavia* | Root and Soil |
| dFASV19 | Ascomycota | *Sordariomycetes* | Root and Soil |
| dFASV20 | Ascomycota | *Setophoma* | Root and Soil |
| dFASV21 | Ascomycota | *Talaromyces* | Root and Soil |
| dFASV22 | Ascomycota | *Trichoderma* | Root |
| dFASV23 | Ascomycota | *Humicola* | Root and Soil |
| dFASV24 | Ascomycota | *Epicoccum* | Root |
| dFASV25 | Ascomycota | *Penicillium* | Root and Soil |
| dFASV26 | Ascomycota | *Penicillium* | Root and Soil |
| dFASV27 | Ascomycota | *Pseudopithomyces* | Root |
| dFASV28 | Ascomycota | *Scytalidium* | Root |
| dFASV29 | Ascomycota | *Trichurus* | Root and Soil |
| dFASV30 | Ascomycota | *Sarocladium* | Root and Soil |
| dFASV31 | Ascomycota | *Aspergillus* | Root and Soil |
| dFASV32 | Ascomycota | *Geomyces* | Root and Soil |
| dFASV33 | Ascomycota | *Blumeria* | Root and Soil |
| dFASV34 | Ascomycota | *Cyphellophora* | Root |
| dFASV35 | Ascomycota | *Fusarium* | Root |
| dFASV36 | Ascomycota | *Ascomycete* | Root and Soil |
| dFASV37 | Ascomycota | *Pyrenophora* | Root |
| dFASV38 | Ascomycota | *Penicillium* | Root and Soil |
| dFASV39 | Ascomycota | *Gibellulopsis* | Root |
| dFASV40 | Ascomycota | *Fusarium* | Root |
| dFASV41 | Ascomycota | *Conlarium* | Root |
| dFASV42 | Ascomycota | *Metarhizium* | Root and Soil |
| dFASV43 | Ascomycota | *Dothideomycetes* | Root |
| dFASV44 | Ascomycota | *Paecilomyces* | Root and Soil |
| dFASV45 | Ascomycota | *Ochroconis* | Root |
| dFASV46 | Ascomycota | *Fusarium* | Root |
| dFASV47 | Ascomycota | *Uncultured Ascomycota* | Root |
| dFASV48 | Ascomycota | *Uncultured Ascomycota* | Root |
| dFASV49 | Ascomycota | *Nectria* | Root |
| dFASV50 | Ascomycota | *Uncultured Ascomycota* | Root |
| dFASV51 | Ascomycota | *Trichoderma* | Root and Soil |
| dFASV52 | Ascomycota | *Fusarium* | Root |
| dFASV53 | Ascomycota | *Trichothecium* | Root |
| dFASV54 | Ascomycota | *Tetraplosphaer* | Root |
| dFASV55 | Ascomycota | *Uncultured Ascomycota* | Soil |
| dFASV56 | Ascomycota | *Uncultured Ascomycota* | Soil |
| dFASV57 | Ascomycota | *Fusarium* | Soil |
| dFASV58 | Ascomycota | *Fusarium* | Soil |
| dFASV59 | Ascomycota | *Chaetosphaeria* | Soil |
| dFASV60 | Ascomycota | *Acremonium* | Soil |
| dFASV61 | Ascomycota | *Penicillium* | Soil |
| dFASV62 | Ascomycota | *Uncultured Ascomycota* | Soil |
| dFASV63 | Ascomycota | *Aspergillus* | Soil |
| dFASV64 | Ascomycota | *Mucor* | Soil |
| dFASV65 | Ascomycota | *Pseudogymnoasc* | Soil |
| dFASV66 | Ascomycota | *Absidia* | Soil |
| dFASV67 | Ascomycota | *Mucor* | Soil |
| dFASV68 | Ascomycota | *Chloridium* | Soil |
| dFASV69 | Ascomycota | *Uncultured Ascomycota* | Soil |
| dFASV70 | Ascomycota | *Uncultured Ascomycota* | Soil |
| dFASV71 | Basidiomycota | *Uncultured Basidiomycota* | Soil |
| dFASV72 | Ascomycota | *Aspergillus* | Soil |
| dFASV73 | Ascomycota | *Uncultured Ascomycota* | Soil |
| dFASV74 | Ascomycota | *Penicillium* | Soil |
| dFASV75 | Ascomycota | *Penicillium* | Soil |
| dFASV76 | Basidiomycota | *Uncultured Basidiomycota* | Soil |
| dFASV77 | Ascomycota | *Lecythophora* | Soil |
| dFASV78 | Ascomycota | *Chaetomium* | Soil |
| dFASV79 | Ascomycota | *Achaetomium* | Soil |

**Supplementary Table 6:** Differentially abundant bacterial ASVs (dBASVs), their plant compartments, phylum, and genus

| **ID** | **Phylum** | ***Genus*** | **Compartment** |
| --- | --- | --- | --- |
| dBASV1 | Firmicutes | *Neobacillus* | Root and Soil |
| dBASV2 | Deinococcus-Thermus | *Deinococcus* | Root and Soil |
| dBASV3 | Firmicutes | *Bacillus* | Root and Soil |
| dBASV4 | Actinobacteria | *Streptomyces* | Leaf |
| dBASV5 | Actinobacteria | *Solirubrobacter* | Leaf |
| dBASV6 | Actinobacteria | *Catellatospora* | Leaf |
| dBASV7 | Actinobacteria | *Humibacter* | Leaf |
| dBASV8 | Actinobacteria | *Humibacter* | Leaf |
| dBASV9 | Firmicutes | *Staphylococcus* | Leaf |
| dBASV10 | Proteobacteria | *Xanthomonas* | Leaf |
| dBASV11 | Proteobacteria | *Xanthomonas* | Leaf |
| dBASV12 | Proteobacteria | *Unculture Chromatiales* | Leaf |
| dBASV13 | Proteobacteria | *Uncultured Halomonadaceae* | Leaf |
| dBASV14 | Firmicutes | *Staphylococcus* | Leaf |
| dBASV15 | Firmicutes | *Staphylococcus* | Leaf |
| dBASV16 | Acidobaceteria | *uncultured Acidobacteria* | Root |
| dBASV17 | Actinobacteria | *Streptacidiphilus* | Root |
| dBASV18 | Planctomycetes | *Singulisphaera* | Root |
| dBASV19 | Actinobacteria | *Streptomyces* | Root |
| dBASV20 | Proteobacteria | *Unculture Acetobacteraceae* | Root |
| dBASV21 | Actinobacteria | *Streptomyces* | Root |
| dBASV22 | Actinobacteria | *Streptomyces* | Root |
| dBASV23 | Proteobacteria | *Amorphus* | Root |
| dBASV24 | Actinobacteria | *Verrucosispora* | Root |
| dBASV25 | Proteobacteria | *Rhizobium* | Root |
| dBASV26 | Proteobacteria | *Amphorus* | Root |
| dBASV27 | Proteobacteria | *Bradyrhizobium* | Root |
| dBASV28 | Proteobacteria | *Sphingomonas* | Root |
| dBASV29 | Bacteroidetes | *Hymenobacter* | Root |
| dBASV30 | Firmicutes | *Bacillus* | Root |
| dBASV31 | Proteobacteria | *Phyllobacteriaceae_unclassified* | Root |
| dBASV32 | Proteobacteria | *Phyllobacteriaceae_unclassified* | Root |
| dBASV33 | Actinobacteria | *Frankia* | Root |
| dBASV34 | Actinobacteria | *Streptomyces* | Root |
| dBASV35 | Actinobacteria | *Actinokineospora* | Root |
| dBASV36 | Uidentified | *uncultured bacteria* | Root |
| dBASV37 | Firmicutes | *Paenibacillus* | Root |
| dBASV38 | Proteobacteria | *Sphingomonas* | Root |
| dBASV39 | Proteobacteria | *uncultured Proteobacteria* | Root |
| dBASV40 | Firmicutes | *Bacillus* | Root |
| dBASV41 | Proteobacteria | *Pseudomonas* | Root |
| dBASV42 | Unidentified | *uncultured bacteria* | Root |
| dBASV43 | Unidentified | *uncultured bacteria* | Soil |
| dBASV44 | Actinobacteria | *Pseudonocardia* | Soil |
| dBASV45 | Actinobacteria | *Jatrophihabitans* | Soil |
| dBASV46 | Actinobacteria | *Rathayibacter* | Soil |
| dBASV47 | Unidentified | *uncultured bacteria* | Soil |
| dBASV48 | Actinobacteria | *Skermania* | Soil |
| dBASV49 | Unidentified | *uncultured bacteria* | Soil |
| dBASV50 | Chloroflexi | *uncultured Chloroflexi* | Soil |
| dBASV51 | Chloroflexi | *uncultured Chloroflexi* | Soil |
| dBASV52 | Actinobacteria | *Frankia* | Soil |
| dBASV53 | Proteobacteria | *Pseudomonas* | Soil |
| dBASV54 | Proteobacteria | *Rhizobium* | Soil |
| dBASV55 | Proteobacteria | *Desulforapulum* | Soil |
| dBASV56 | Proteobacteria | *Xanthobacter* | Soil |
| dBASV57 | Proteobacteria | *Dokdonella* | Soil |
| dBASV58 | Proteobacteria | *Rhizobium* | Soil |
| dBASV59 | Actinobacteria | *Gaiella* | Soil |
| dBASV60 | Proteobacteria | *Sphingomonas* | Soil |
| dBASV61 | Actinobacteria | *Frankia* | Soil |
| dBASV62 | Proteobacteria | *Acidovorax* | Soil |
| dBASV63 | Actinobacteria | *Nocardioides* | Soil |
| dBASV64 | Proteobacteria | *Pyruvatibacter* | Soil |
| dBASV65 | Proteobacteria | *Phreatobacter* | Soil |
| dBASV66 | Proteobacteria | *Methylobacterium* | Soil |
| dBASV67 | Proteobacteria | *Bradyrhizobium* | Soil |
| dBASV68 | Proteobacteria | *Mesorhizobium* | Soil |
| dBASV69 | Proteobacteria | *Sinorhizobium* | Soil |
| dBASV70 | Actinobacteria | *Nocardioides* | Soil |
| dBASV71 | Firmicutes | *Bacillus* | Soil |

**Supplementary Table 7:** Comparison between host phylogenetic distances and Bray-Curtis dissimilarity of their microbiomes from glasshouse experiment

| Comparison | Fungi_Bray_Curtis | Host_phylogeny | Bacteria Bray_Curtis |
| --- | --- | --- | --- |
| Pha/Luc | 0.741231 | 0.37 | 0.952708 |
| Pha/Rho | 0.577843 | 0.046 | 0.777207 |
| Pha/Rye | 0.636959 | 0.025 | 0.807435 |
| Pha/Dig | 0.616588 | 0.048 | 0.796276 |
| Luc/Rho | 0.748137 | 0.363 | 0.942205 |
| Luc/Rye | 0.791131 | 0.373 | 0.953195 |
| Luc/Dig | 0.696861 | 0.365 | 0.926341 |
| Rho/Rye | 0.577157 | 0.049 | 0.800882 |
| Rho/Dig | 0.519701 | 0.04 | 0.747516 |
| Rye/Dig | 0.632798 | 0.051 | 0.841235 |

**Supplementary Table 8:** Comparison between host phylogenetic distances and Bray-Curtis dissimilarity of their microbiomes from additional data

| Comparison | Bray_Curtis | Host_Phylogeny |
| --- | --- | --- |
| Wheat/Barley | 0.53322 | 0.015 |
| Wheat/Maize | 0.922421 | 0.143 |
| Wheat/Luc | 0.977009 | 0.382 |
| Wheat/Euc | 0.985427 | 0.218 |
| Wheat/Rho | 0.981351 | 0.136 |
| Wheat/Wal | 0.977082 | 0.147 |
| Wheat/Cot | 0.988434 | 0.234 |
| Barley/Maize | 0.921755 | 0.142 |
| Barley/Euc | 0.991148 | 0.204 |
| Barley/Rho | 0.99039 | 0.143 |
| Barley/Wal | 0.989373 | 0.148 |
| Barley/Cot | 0.992963 | 0.241 |
| Maize/Euc | 0.991559 | 0.3 |
| Maize/Rho | 0.915426 | 0.041 |
| Maize/Wal | 0.897281 | 0.045 |
| Maize/Cot | 0.897281 | 0.23 |
| Euc/Rho | 0.960857 | 0.201 |
| Euc/Wal | 0.985282 | 0.205 |
| Euc/Cot | 0.509986 | 0.146 |
| Rho/Wal | 0.758791 | 0.046 |
| Rho/Cot | 0.99039 | 0.231 |
| Wal/Cot | 0.989373 | 0.235 |
| Kang/Cot | 0.9862 | 0.338 |
| Kang/Euc | 0.9531 | 0.27 |
| Kang/Rho | 0.731 | 0.167 |
| Kang/Maize | 0.701 | 0.165 |
| Kang/Wal | 0.5535 | 0.155 |
| Kang/Barley | 0.784 | 0.165 |
